# AGO5 restricts virus vertical transmission in plant gametophytes

**DOI:** 10.1101/2025.11.28.691182

**Authors:** Gesa Hoffmann, Sri Pravallika Sadhu, Gabriele Bradamante, Juan C. Diez-Marulanda, Antonia Proschwitz, Tobias Wegscheider, Ilayda Turhan, Heinrich Bente, Ruben Gutzat, Marco Incarbone

## Abstract

The propagation of a viral infection from a host parent to its progeny is known as vertical transmission, or seed transmission in plants. It allows viral infections to rapidly spread locally via pollen and worldwide through seeds. To be vertically transmitted to each progeny, a virus must pass through the tight bottleneck of at least one cell per parent – the gametes. Therefore, stopping infection during sexual reproduction is of vital importance to generate healthy offspring. Accordingly, vertical transmission of plant viruses often occurs at very low rates, if at all, suggesting the existence of highly effective – yet unknown – antiviral defenses in pre-meiotic cells, gametes and/or embryos. In this study, we show that AGO5, an RNA interference factor expressed specifically in shoot apical meristem stem cells and the germline of *Arabidopsis thaliana*, drastically reduces the vertical transmission of Turnip yellow mosaic virus (TYMV). Through a series of controlled pollination experiments leveraging different zygosity of *ago5* knock-out, cell type-specific rescue of *ago5* and TYMV detection in whole-mount reproductive tissues, we provide evidence that AGO5 acts in pollen and sperm cells to restrict virus transmission to progeny. We further show that triggering antiviral RNA interference specifically in sperm cells leads to a significant reduction in TYMV vertical transmission. In summary, this study provides the first description of a gamete-specific antiviral defense mechanism restricting virus vertical transmission, paving the way for new strategies to prevent the spread of pollen- and seed-borne viral epidemics.

## MAIN TEXT

Viruses are obligate intracellular parasites that require living hosts and need to be transmitted to new ones to ensure their continued existence. Viral host-to-host transmission can essentially occur in two directions: vertically between a host and its progeny or horizontally between hosts that do not necessarily share parent/progeny relations. Horizontal virus transmission mechanisms have been extensively investigated in virology, yet the molecular events underlying vertical transmission (VT) and the defenses deployed by the host to prevent it remain very poorly understood. In sexually reproducing organisms, *bona fide* VT occurs through the germline, the cells passing genetic information to the progeny. Infection of progeny by maternal tissues after fertilization, as well as transmission mediated by the gamete companion cells such as the vegetative cell in pollen, can also be considered a form of VT, albeit not by cells passing on genetic inheritance. When discussing VT, an important distinction must be made between (i) viral and virus-like genetic information within the host genome, which is vertically transmitted by default through host DNA replication, and (ii) extra-genomic viruses which must ensure the inheritance of their genetic information independently of host genome replication (*1*). Plant viruses essentially belong to the second category, since infectious endogenous viral elements in plant genomes are not common (*2, 3*) and the majority of described virus species have RNA genomes (*4, 5*).

Plant viruses constitute a prime threat to crop production worldwide and vertical (or seed) transmission of viral diseases has the potential to cause their rapid local and global diffusion due to the high mobility of pollen, carrying the male germline, and the worldwide exchange of seed stocks (*6-8*). Interestingly though, many viral infections result in little to no VT to progeny despite efficient infection of the parent, suggesting that plants deploy potent transgenerational antiviral defenses to ensure a healthy offspring (*1*). Little is known about the cell types and developmental stages through which VT can occur or be blocked, although the accepted model proposes two non-mutually exclusive options: (i) the germline-precursor stem cells before meiosis and gametophyte formation or (ii) the developing embryo after fertilization (*9*). It is likely that the well-established ability of stem cells within the shoot apical meristem (SAM) to remain free of viral infection can play a role in limiting virus transmission through the subsequently developing germline, although direct evidence of this is lacking. Equally, our knowledge on the molecular mechanisms regulating VT remains very limited: RNA interference (RNAi) and its suppression by viruses have been shown to be major determinants (*10-13*) along with the speed of infection spread within the parent plants (*14*). Finally, viruses that produce cryptic/asymptomatic infections can transmit vertically at very high rates (*15*), broadly suggesting an inverse relation between pathogenicity in the parent and efficient transmission to the progeny (*16, 17*).

RNAi is one of the main and most conserved antiviral pathways in plants (*3*). It is initiated by viral double-stranded RNA, an inevitable intermediate of RNA virus replication and a hallmark of infection, which is processed by endoribonucleases of the dicer family (Dicer-Like - DCL) into 21-22nt-long virus-derived small interfering RNA (vsiRNA). These are loaded into Argonaute (AGO) proteins to mediate sequence-specific recognition and degradation/neutralization of viral RNA (*3*). Subsequently, viruses have necessarily evolved numerous strategies and effectors to suppress or evade RNAi (*18*). RNAi and its amplification by host-encoded RNA-dependent RNA polymerases (RDR) play a key role in SAM stem cell immunity and the fertility of infected *A. thaliana* plants (*19, 20*). Knock-out of DCL genes in parent plants increases rates of virus VT (*13*), possibly a consequence of greatly increased virus titer in the parents. AGO5 is one of ten *A. thaliana* AGO proteins, known to have antiviral function (*21-23*) and, compellingly, to be expressed specifically in germline-precursor SAM stem cells and gametes (*24, 25*). Using a mutant of Cucumber mosaic virus (CMV) compromised in its RNAi suppression ability, it has been shown that AGO5 can limit VT but only when *AGO1* and *AGO2* genes are knocked out, suggesting that AGO5 can serve as last-resort backup defense against a recombinant RNAi-hypersensitive virus (*13*). Given these observations, we wondered whether AGO5 could play a role in stem cell and germline antiviral defense, thereby limiting VT of a wild-type, uncompromised virus. But most importantly, we wondered in which cells and stages of development virus VT could be blocked, a fundamental aspect that remains unknown and could prime the development of targeted tools to tackle virus transmission through seed. To investigate this, we used the *A. thaliana*/TYMV (Turnip yellow mosaic virus) pathosystem, which we developed as a model for the systematic investigation of VT in plants. TYMV has a (+)ssRNA genome, infects several Brassicaceae species (*26-30*) and is vertically transmitted at high rates in *A. thaliana* (Arabidopsis hereafter) (*31*).

To investigate whether VT of TYMV is the result of gamete infection, we studied the localization of two TYMV isolates in Arabidopsis meristems and reproductive tissues, first using RNA *in situ* hybridization (ISH) on slices of embedded tissue, then on a larger scale by whole-mount single molecule Fluorescent *In Situ* Hybridization (FISH) (*32*). We have previously shown by ISH that TYMV is excluded from the SAM (*20*) and could confirm these results by FISH (**Fig. 1A**; **Fig. S1A**,**B**). This indicates that TYMV enters the germline at a later developmental stage, but presumably before gametophytes lose symplastic connection to the surrounding tissues. In the gynoecium, TYMV RNA is detectable in the valves and septum, as well as inside ovules (**Fig. 1B,C**; **Fig. S2**). In developing seeds of infected plants 48 hours after pollination with healthy pollen, we observed three different infection patterns: while the inner and outer integuments were fully infected, the embryo and endosperm could either (i) both be virus-free (**Fig. 1D, Fig. S3**), (ii) the endosperm infected but not the embryo (**Fig. 1E, Fig. S3**) or both be infected (**Fig. 1F, Fig. S3**). This indicates that both egg cell and central cell can be infected by TYMV prior to fertilization, but this is not always the case despite pervasive infection of the surrounding sporophytic tissues. Whether TYMV is capable of moving between integuments and endosperm or invade the embryo through the suspensor (as shown for Pea seedborne mosaic virus (*33*)), remains to be determined. In addition to the female reproductive tissues, TYMV accumulates in the stamen (**Fig. 1G, Fig. S4A**) and in a number of pollen grains where viral RNA can be detected in the vegetative cell, as well as the sperm cells (**Fig. 1H, Figs. S4B, S5**). By pollinating non-infected plants with pollen from TYMV-infected plants, we detected TYMV-positive pollen tubes 24 h after pollination (**Fig. 1I, Fig. S6A**) and virus accumulation in early endosperm at 24 h (**Fig. 1J, Fig. S6B**) and in zygote and endosperm at 48 h (**Fig. 1K, Fig. S7**) after pollination, clearly indicating that TYMV is vertically transmitted through male gametophytes in Arabidopsis. Interestingly, just like in female transmission, male-transmitted TYMV RNA can be absent from the early embryo, but present in the endosperm, in line with localization in the pollen vegetative cell and/or only one of the sperm cells in a pollen grain being infectious (**Fig. 1L**). These results clearly show that TYMV is capable of infecting both male and female reproductive organs, but also that viral RNA can be excluded from individual gametophytes within the same flower. The effects of TYMV infection on the viability and development of each cell type, as well as their link to VT remains to be determined.

**Figure 1:**
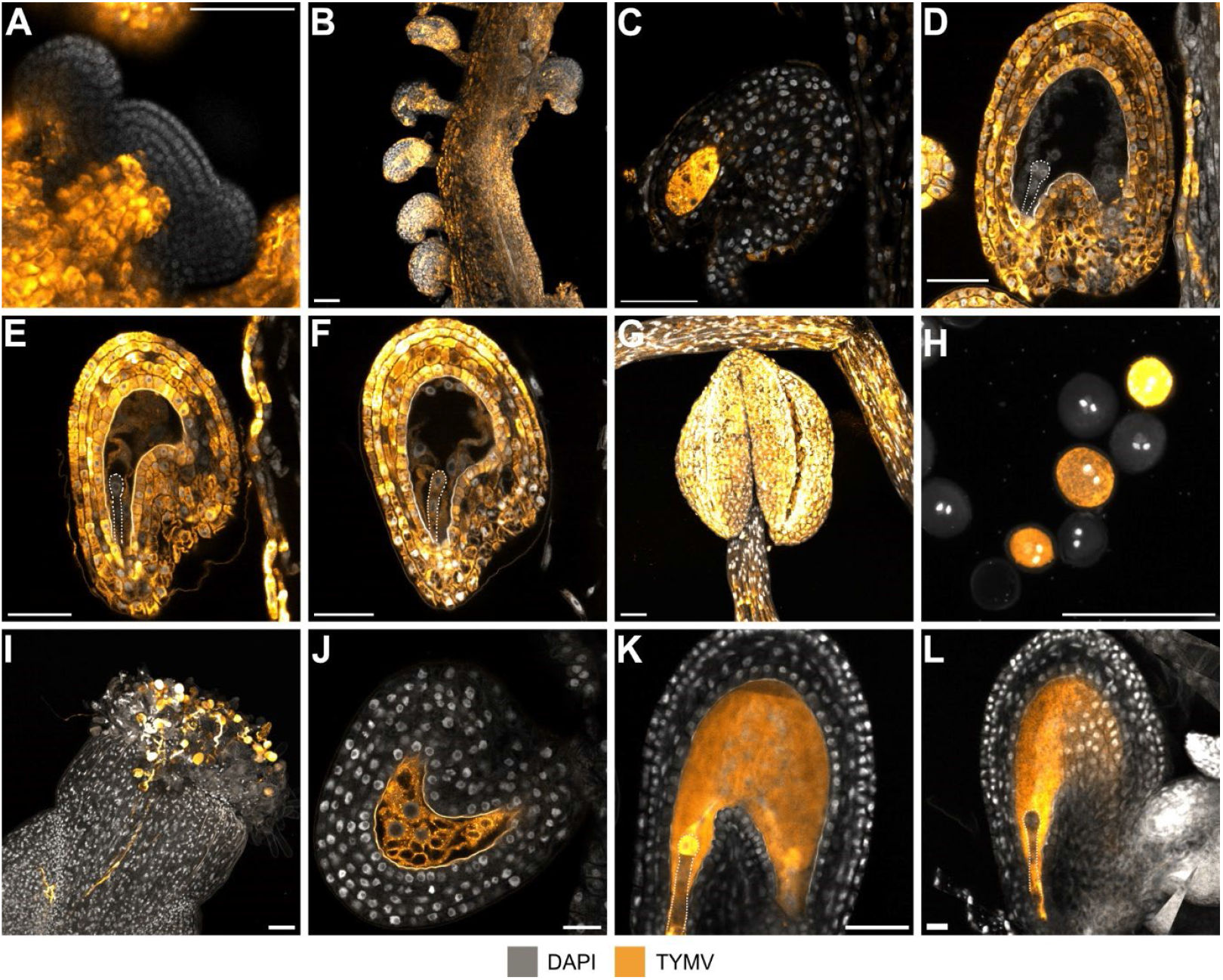
Detection of TYMV in germline and zygote by whole-mount FISH in Arabidopsis reproductive tissues. **(A)** TYMV is excluded from the shoot apical meristem. **(B)** TYMV fully infects gynoecia and **(C)** can be detected at the micropylar pole of the female gametophyte. **(D-F)** Developing seeds from systemically infected maternal plants 48 hours after pollination with healthy pollen show three TYMV infection patterns in the integuments, endosperm and embryo. **(G)** TYMV infects anthers and **(H)** is either fully absent or detected in vegetative and sperm cell cytoplasm in mature pollen grains. **(I**,**J)** 24 hours after pollination of healthy mother plants with pollen from TYMV-infected plants, viral RNA is detected in **(I)** pollen grains on stigmas and pollen tubes, leading to **(J)** endosperm infection. **(K)** 48 hours after pollination TYMV can infect developing embryos and endosperm, although **(L)** some seeds show positive signal only in the endosperm. Images show merge of TYMV FISH channel (pseudo-colored in LUT "Orange Hot”) and DAPI (greyscale). Scale bars = 50 µm. B,G,H and I are max projections. Embryos are outlined in white dotted lines for clarity. Further supporting images available in Supplementary Figures S1-S7.

We assessed TYMV VT rates through self-fertilization of infected parents in a panel of wild-type (WT) and mutants in germline-specific AGO genes: AGO5 and AGO9. We used two TYMV isolates that share 97.96% sequence identity but differ in their symptomology and disease severity. TYMV-S (severe) causes a strong reduction in pollen production and overall fertility of infected plants, while TYMV-M (mild) showed less drastic effects on pollen production and fertility compared to TYMV-S (**Fig. S8A-D**). Most importantly, successful VT of either isolate induces visible yellowing of leaves in emerging progeny seedlings, which corresponds to the presence of virus, making them easily distinguishable from virus-free plants by eye (**Fig. 2A, Fig. S8E**). This scalable system allows rapid and reliable quantification of VT in large numbers of parent plants and progeny seedlings, a crucial requirement to overcome the substantial variations between individual parents. Two independent *ago5* T-DNA insertion knock-out mutant alleles, *ago5-1* and *ago5-6*, showed significantly higher VT rates of both isolates compared to WT (**Fig. 2B,C**). Conversely, knock-out of *AGO9*, a nuclear-localized AGO sharing similar expression patterns as *AGO5* (*24*), produced no significant differences in VT (**Fig. 2B,C**). Three independent transgenic lines expressing a MYC-tagged AGO5 under the native *pAGO5* promoter in the *ago5* background could restore VT to WT levels (**Fig. 2D,E**). Interestingly, *ago5* mutants showed no plant-wide differences in symptoms (**Fig. S8A**,**B**) or viral RNA accumulation in inflorescences (**Fig. S8F**). It is especially important to note that the previously described meristematic virus exclusion zone for TYMV consisting of the top 2-3 cell layers of the flowering SAM (*20*) was not abrogated in *ago5* mutants (**Fig. S1C**,**D**), as was reported for other RNAi mutants (*20*), suggesting that increased VT is not necessarily a consequence of increased SAM stem cell infection. Taken together, these results clearly indicate that AGO5 plays a cell type-specific antiviral role in the restriction of VT. We next wondered whether AGO5 restricts virus VT in the gametophytes, the crucial unexplored bottleneck in transgenerational infection. We focused subsequent experiments on pollen transmission of TYMV because (i) our FISH experiments revealed prominent viral RNA accumulation in pollen, suggesting that pollen can transmit TYMV and (ii) transmission via pollen can be used as an accurate sensor of gametophyte- and gamete-mediated VT, in contrast to egg-mediated transmission for which infection from the surrounding maternal tissues cannot be excluded. First, we tested whether AGO5 restricts VT via pollen, in accordance with its expression in sperm cells (*24*). Crossing experiments with infected male and non-infected female parents showed that AGO5 strongly reduces virus VT via pollen (**Fig. 3A**). We then leveraged this system to assess whether AGO5 restricts VT pre-meiosis in the sporophyte, post-meiosis in the gametophytes, or post-fertilization in the zygote and embryo. Hereafter, all experiments are performed with the TYMV-M isolate. First, we probed for a potential pre-meiosis effect of AGO5 on VT. It is generally assumed that, due to the establishment of symplastic isolation of the male gametophyte early in development, virus infection of the gametophyte precursor cells must occur before meiosis (*34, 35*). In accordance, pollen tetrads of infected *qrt1-4* parent plants were either fully infected or free of virus. *qrt1-4* mutant plants show a defect in pollen grain separation after microsporogenesis (*36*) leading to developmentally linked tetrad pollen that share a common microspore mother cell precursor, strongly supporting that TYMV infects pollen precursors before meiosis (**Fig. 3B; Fig. S9**). We therefore reasoned that if AGO5 exerts antiviral activity and restricts virus establishment/proliferation in the germline precursor cells before meiosis, *ago5* plants should show an increased number of infected gametophytes compared to WT. Large-scale FISH quantification of TYMV-positive pollen grains revealed that, on the contrary, the number of TYMV-positive pollen grains were not significantly different between WT and *ago5* (**Fig. 3C; Fig. S10**), and viral RNA accumulation in pollen was comparable **(Fig. S11A)**. This suggests that AGO5 exerts its role to limit VT later during development and is not able to clear TYMV from infected pollen grains. Together, these results indicate that AGO5 does not significantly affect VT pre-meiotically.

**Figure 2:**
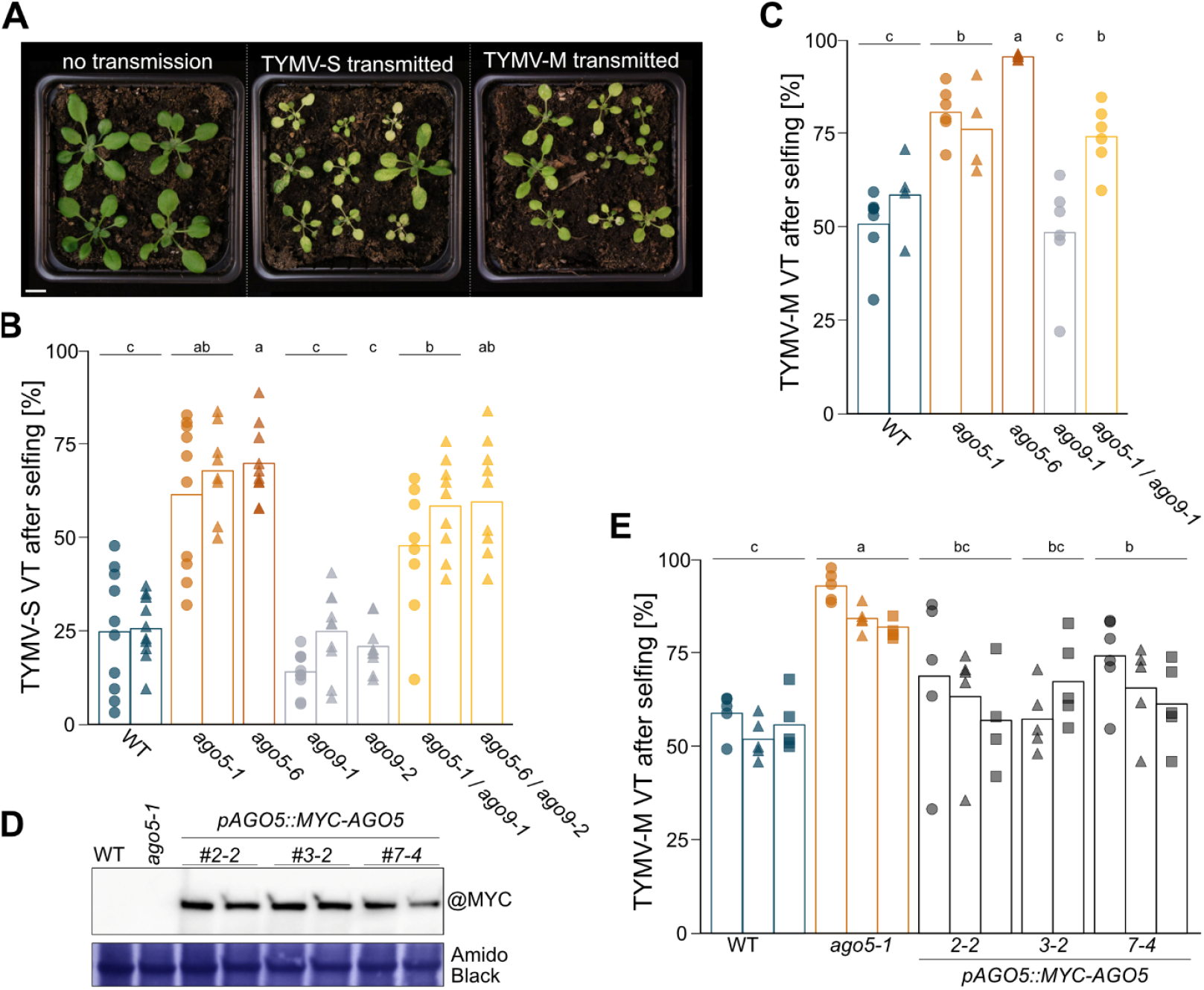
Arabidopsis AGO5 restricts TYMV vertical transmission. **(A)** Representative image of progeny from infected WT plants showing healthy and infected seedlings 21 days after sowing (scale bar = 1 cm). **(B)** VT rates [%] of TYMV-S in indicated genotypes after selfing. Single data points refer to VT from individual parent plants, shape of data points to independent infection experiments. Seedlings counted (sc): 19164. **(C)** VT rates [%] of TYMV-M in indicated genotypes after selfing. sc: 9437. **(D)** Detection of MYC:AGO5 expression in three independent transgenic lines of *pAGO5::MYC:AGO5*/*ago5-1* inflorescence tissue, in biological duplicates. **(E)** VT rates [%] of TYMV-M in *pAGO5::MYC:AGO5*/*ago5-1* complementation lines. sc: 18690. In B, C and E statistical significance was determined by one-way ANOVA coupled with Tukey’s HSD test (*α* = 0.05), letters indicate statistical groups. Vertical bars indicate average of datapoints from single parent plants depicted by shapes. B, C and E each represent data from independent sets of experiments.

**Figure 3:**
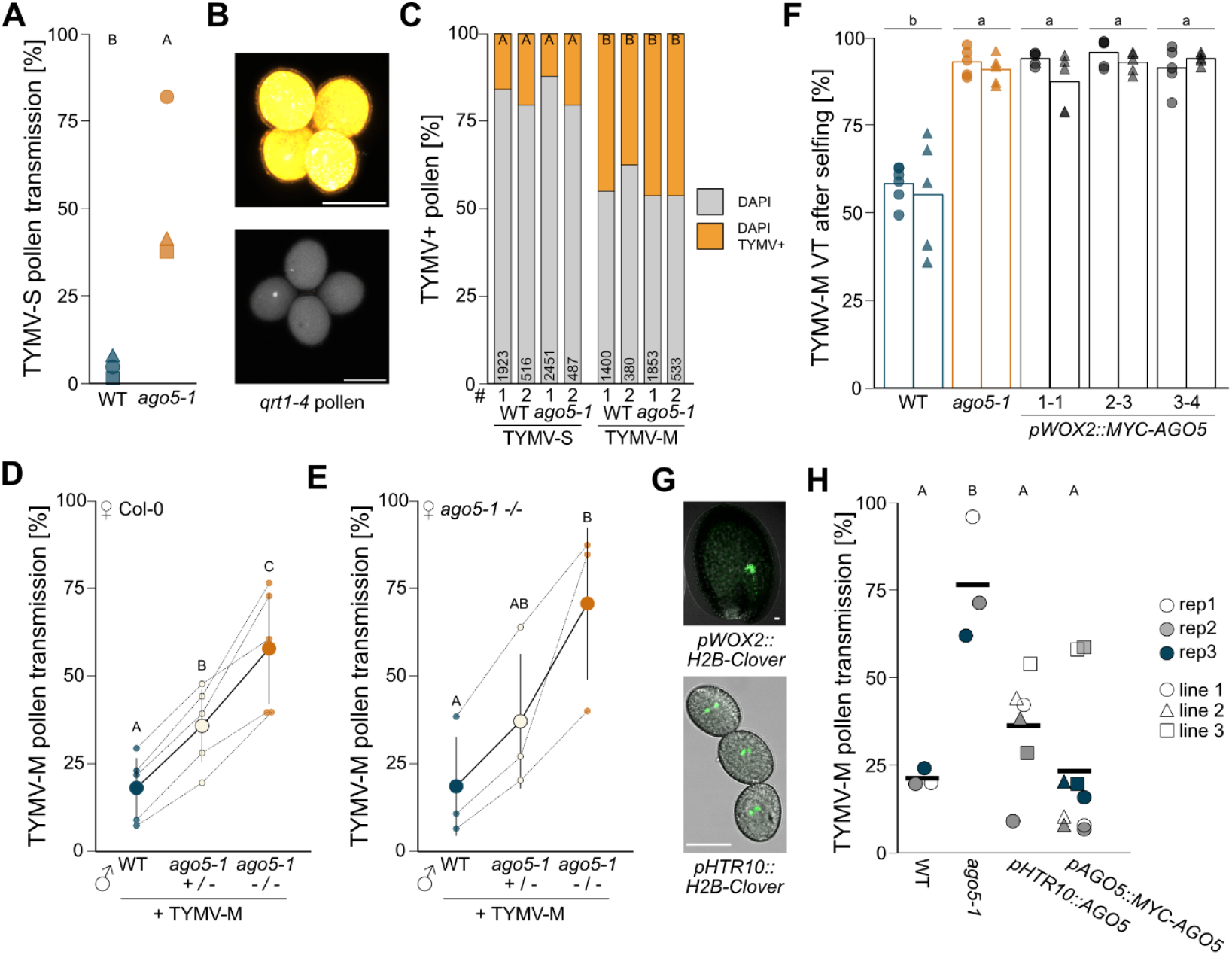
AGO5 restricts TYMV VT via pollen after meiosis but before fertilization, in gametes and gametophytes. **(A)** VT via pollen rate [%] of TYMV-S after pollination of non-infected WT mother plants with pollen from TYMV-infected WT or *ago5-1*. Shape of data points refer to pollinations from chronologically separated infection experiments. Seedlings counted (sc): 3495. **(B)** Max projection of FISH detection of TYMV genomic RNA in pollen tetrads from TYMV-M infected *qrt1-4* plants (scale bar = 20 µm). **(C)** Fraction of pollen with positive FISH signal for TYMV in indicated genotypes. # refers to the experiment. Numbers in bars indicate n of pollen grains counted. **(D)** VT via pollen rate [%] of TYMV-M after pollination of non-infected WT mother plants with pollen from TYMV-infected WT (+/+), *ago5-1* heterozygous (+/-) or *ago5-1* homozygous (-/-) mutants. Small dots indicate individual pollination experiments, large dots indicate their averages. Datapoints from the same experiment are connected by dashed lines. sc: 7503. **(E)** As in (D) but using non-infected *ago5-1* mutants as mother plants. sc: 2995. **(F)** VT rates [%] of TYMV-M after selfing in three independent transgenic lines of *pWOX2::MYC:AGO5*/*ago5-1*. Single data points refer to individual parent plants, shape of data points to independent infection experiments. Vertical bars indicate average of datapoints from single parent plants depicted by shapes. sc: 11825. **(G)** Laser confocal microscopy images of *pWOX2::H2B:Clover*/WT and *pHTR10::H2B:Clover*/WT reporter lines showing specific expression in the embryo (top) and in the sperm cells of mature pollen grains (bottom), respectively. Scale bar = 20 µm. **(H)** VT via pollen rate [%] of TYMV-M in three independent transgenic lines of *pAGO5::MYC:AGO5*/*ago5-1* and *pHTR10::AGO5*/*ago5-1*. Color of data points refer to pollinations from chronologically separated infection experiments, shapes indicate independent lines. sc: 6127. In A, C, D, E and H statistical comparison was done per genotype by applying generalized linear models (GLMs) with a quasibinomial distribution, statistical groups are plotted above in capital letters (p < 0.05). In F statistical significance was determined by one-way ANOVA coupled with Tukey’s HSD test (*α* = 0.05), lower case letters indicate statistical groups.

To address whether AGO5 restricts VT post-meiosis, we performed controlled crosses with either WT, *ago5*+/-heterozygous, or *ago5*-/-homozygous mutants as infected pollen donors, while the receiving non-infected mothers were WT. In *ago5*+/-plants all pre-meiotic cells express the *AGO5* gene, while only half the gametophytes express *AGO5* due to segregation of the mutant allele during meiosis. Therefore, excluding possible dosage effects pre-meiosis, if AGO5 blocks VT after meiosis, the VT levels from an *ago5*+/-virus donor are expected to be at 50% between those from WT and *ago5*-/-. Accordingly, the crossing experiments showed that on average *ago5*+/-yielded VT levels at 47.1% between WT and *ago5*-/-(**Fig. 3D**), strongly supporting the hypothesis that AGO5 limits VT after meiosis in male gametophytes. Interestingly, progeny of selfed *ago5+/-*also exhibited an intermediate transmission rate (**Fig. S11B**), suggesting that AGO5 might work similarly in the female germline. AGO5 could also limit VT after fertilization has occurred. In this case, the genotype of the non-infected parent introducing the mutant allele after fertilization would affect VT rates due to the resulting 50% *ago5-/-*progeny. We therefore repeated the crosses as above (**Fig. 3D**) but replacing WT with *ago5*-/-as non-infected mother plants. The *ago5-/-*mutation on the maternal side had no effect on TYMV transmission rates (**Fig. 3E**) when compared to the crosses with functional AGO5 on the maternal side (**Fig. 3D**), showing that AGO5 does not affect VT post-fertilization in the zygote and embryo. Importantly, this independent set of crosses confirmed the intermediate VT phenotype given by *ago5+/-*pollen and WT mothers, reiterating the post-meiotic effect of AGO5. To corroborate that AGO5 is not involved in the antiviral defense after fertilization, we complemented the *ago5* mutation with MYC:AGO5 under the zygote/embryo-specific promoter *pWOX2* (*37, 38*). In accordance with our findings described above, expression of AGO5 exclusively in embryos did not reduce VT in three independent transgenic lines (**Fig. 3F,G**). These results prompted us to further probe the male gamete-specific role of AGO5 in the reduction of virus VT, as AGO5 and AGO9 are the only AGO genes expressed in sperm cells (*24*). AGO5 complementation lines under the native *pAGO5* promoter reduced TYMV VT via pollen (**Fig. 3H**). We next used the *pHTR10 (39)* promoter to drive *AGO5* expression exclusively in sperm cells of *ago5* mutants (**Fig. 3G**). Crossing experiments with three independent transgenic lines showed that sperm cell-specific expression of *AGO5* substantially reduced VT via pollen (**Fig. 3H**), confirming the post-meiotic and gametic restriction of VT by AGO5. AGO proteins are well known to mediate antiviral activity in plants, as their abrogation leads to increased viral fitness (*40*). AGOs can perform direct antiviral RNAi by loading vsiRNA and targeting viral nucleic acids, as shown with *in vitro* systems (*41*), and can indirectly modulate infection by loading host small RNA and regulating gene expression (*42, 43*). Although the targeting of viral RNA by AGOs remains the most plausible mechanism, the precise contributions of the direct vs. indirect aspects of AGO activity to the limitation of VT *in vivo* remains to be causally and genetically established.

Given the crucial role for antiviral RNAi in gametes uncovered in our study, we wondered if this knowledge could be applied to effectively reduce viral propagation by priming defenses in specific cells at the bottleneck of transmission. To determine whether antiviral RNAi can be primed to reduce TYMV VT, we generated lines expressing an RNA hairpin to trigger production of TYMV-specific siRNA either ubiquitously, in the sperm cells, or in the zygote/embryo. As controls, we generated lines expressing a hairpin that does not share sequence homology with TYMV, derived from the fluorescent protein Scarlet (*20*). Small RNA sequencing from the sperm cell-specific *pHTR10::siTYMV/siScarlet* lines confirmed abundant siRNA production from the respective hairpin construct in pollen (**Fig. 4A, Fig. S12A**). Crossing experiments with these lines revealed that triggering virus-specific RNAi in sperm cells reduced VT via pollen by 52% on average (**Fig. 4B, Fig. S13A**). Surprisingly, expression of the non-homologous small RNAs generally increased VT of TYMV compared to WT (**Fig. 4B**), suggesting that a non-virus-related siRNA burden on the RNAi machinery may favor VT. Conversely, the production of the well-studied viral RNAi suppressor protein P19 (*44*) in sperm cells led to an increase in TYMV pollen transmission of 96% (**Fig. 4B, Fig. S13B**). These findings confirm that sperm-cell specific RNAi can be primed to limit VT of TYMV. As a comparison to assess the effectiveness of cell-type-specific RNAi in reducing VT, we also expressed the hairpin constructs ubiquitously in parental plants. This reduced TYMV VT by 32%-52% (**Fig. 4C, Fig. S13C**), while increasing plant fecundity, an indication of reduced disease (**Fig. S12B)**. Interestingly, zygote- and embryo-specific hairpin expression also reduced VT (**Fig. 4D, Fig. S13D**), while not affecting seed production (**Fig. S12C**), indicating that post-fertilization RNAi can decrease virus transmission, but since AGO5 does not restrict VT after fertilization (**Fig. 3F**), this effect is most likely independent of AGO5. Remarkably, sperm cell-specific hairpin expression proved to be as effective as, or even more effective than, ubiquitous or embryo-specific expression in reducing VT. This finding highlights the pivotal role of gametes in limiting viral spread and opens new avenues for developing cell type-specific strategies to combat pollen- and seed-transmitted viruses.

**Figure 4:**
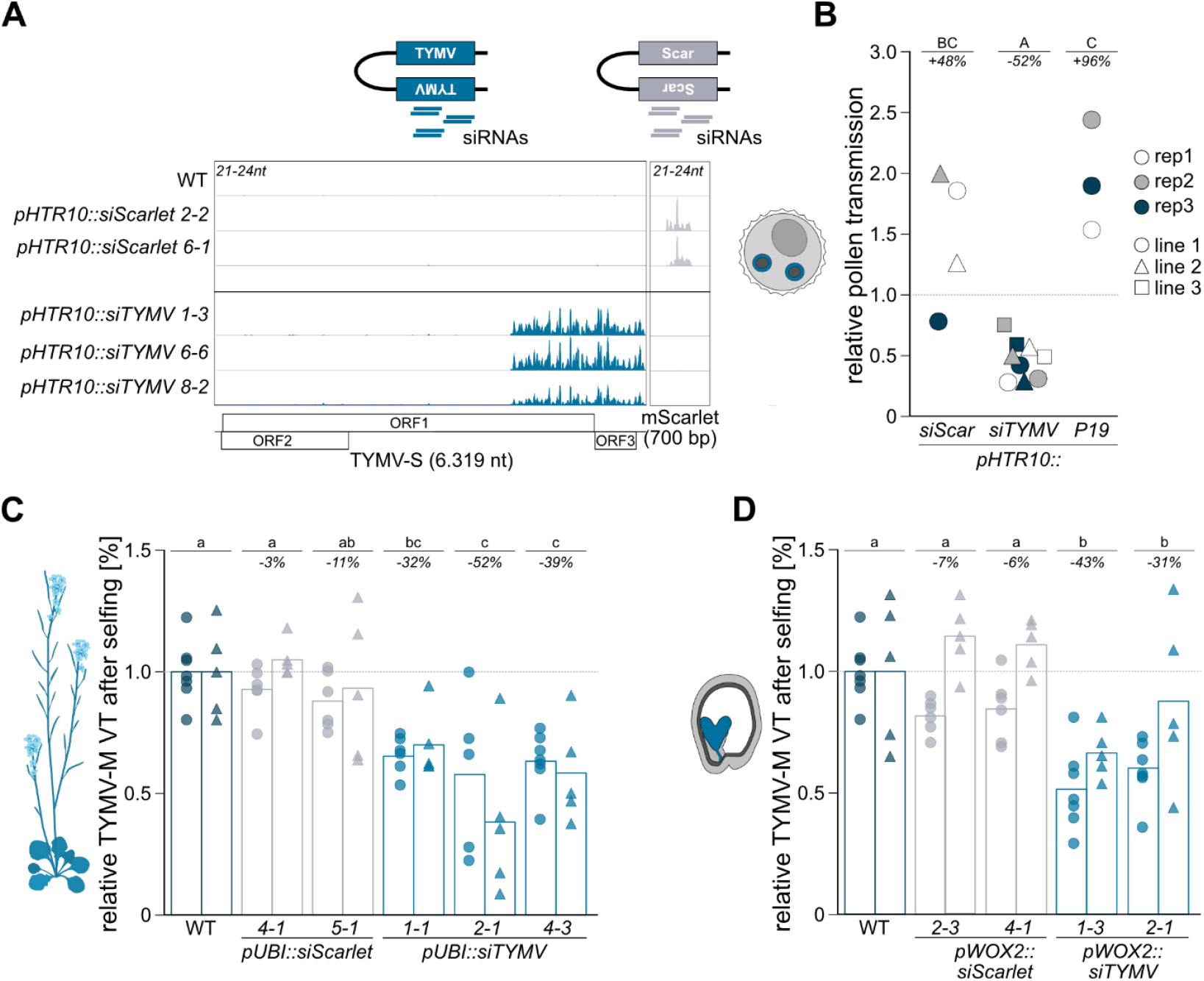
Sperm cell-specific antiviral RNAi can be triggered to reduce TYMV pollen transmission. **(A)** Genome browser shot of 21-24nt-long siRNA populations in mature pollen of three independent lines of *pHTR10::siTYMV*/WT and two of *pHTR10::siScarlet*/WT, showing small RNA reads on the TYMV genome (blue, left) and mScarlet mRNA (grey, right) sequences, respectively. **(B)** Relative VT via pollen of TYMV-M in several transgenic lines of *pHTR10::siScarlet*/WT, *pHTR10::siTYMV*/WT and *pHTR10::P19*/WT. VT via pollen in WT was set to 1 for each replicate (dashed line). Note that WT controls for *pHTR10::siTYMV/Scarlet*/WT constructs replicate 1 and P19 replicate 2 are the same. Color of data points refer to pollinations from chronologically separated infection experiments, shapes indicate independent lines. Seedlings counted (sc): 4516. **(C)** Relative VT of TYMV-M after selfing in several independent transgenic lines of *pUBI::siScarlet*/WT and *pUBI::siTYMV*/WT. Average of individual WT plants was set to 1. Vertical bars indicate average of datapoints from single parent plants depicted by shapes. sc: 19394. **(D)** As in (C) but on *pWOX2::siScarlet*/WT and *pWOX2::siTYMV*/WT. sc: 14288. Note that WT replicate 1 is the same for (C) and (D). Absolute, non-relative VT data available in Fig. S13. In B statistical comparison was done per genotype by applying generalized linear models (GLMs) with a quasibinomial distribution, statistical groups are plotted above in capital letters (*p* < 0.05) Note that WT is statistical group B. In C and D statistical significance was determined by one-way ANOVA coupled with Tukey’s HSD test (*α* = 0.05), lower case letters indicate statistical groups. Representation of plant in C generated with Biorender.

The germline within the gametophytes, where the transmission of pathogens to the next generation involves a single cell per parent, constitutes a critical defensive bottleneck that the host can exploit to curtail inheritance and spread of infection. We have provided compelling evidence that AGO5 serves this precise function, limiting the transmission of a pathogenic RNA virus to the host progeny through antiviral activity specifically in the male gametes and gametophytes, with our complementation experiments indicating germline-specific restriction of transmission. This is the first description of a germline-specific molecular mechanism blocking vertical transmission of an RNA virus and our findings highlight the potential of germline RNAi to challenge the long-range spread of viral epidemics through pollen and seed.

## FUNDING AND ACKNOWLEDGEMENTS

M.I. wishes to acknowledge a Lise Meitner postdoctoral grant from the Austrian Science Fund (FWF M2921) and core research group funding from the Max Planck Society. G.H. acknowledges funding by an EMBO postdoctoral fellowship (ALTF 929-2023) and an MSCA postdoctoral fellowship (101147406 – VirITAS). R.G. gratefully acknowledges financial support from the Austrian Science Fund (I3687). The authors thank Yuki Hamamura, Arun Sampathkumar, Michael Nodine, Vu Nguyen and the MPIMP GreenTeam for assistance with experiments and/or analysis; Frederic Berger, Marion Clavel, Claudia Köhler and Dolf Weijers for insightful suggestions and discussions; Claudia Köhler and Ortrun Mittelsten Scheid for critical feedback on the manuscript. M.I. wishes to especially thank Ortrun Mittelsten Scheid for unwavering support, feedback and mentoring.

## AUTHOR CONTRIBUTIONS

M.I., G.H., G.B. and R.G. conceived and designed the experiments. G.H., A.P., M.I., G.B., R.G. and H.B. performed cloning, infections, crosses, VT rate counting and pollen isolation; G.H., A.P., I.T. and M.I. performed RNA analyses, while J.C.D.M. and A.P. performed protein analysis. G.H. and S.P.S. optimized and performed FISH, while G.B. and T.W. performed ISH. G.H. and M.I. analyzed the data and wrote the manuscript.

## MATERIAL & METHODS

### Plant growth and material

All mutants used in this study were in the background of *Arabidopsis thaliana* accession Columbia-0 (Col-0), which is denoted as wildtype (WT) in this manuscript. Arabidopsis mutant lines *ago5-1 (21), ago9-1 (45), ago9-2 (45)*, and *qrt1-4* (*36*) have been described previously (**Table S2**). *ago5-6* (SAIL_100_C09) was identified as an additional *ago5* mutant. Genotyping was performed by standard PCR of leaf DNA extracts. Double mutants were generated through crossing. Heterozygous *ago5-1*+/-plants for VT assays were obtained by crossing WT and *ago5-1*-/- and using the resulting F1 for the assay. Transgenic lines were generated using flower dip method (*46*); transformants were selected through seed coat fluorescence and propagated to homozygous state. For infection experiments, seeds were stratified overnight in 0.1% agarose and sowed on soil to germinate for one week at 75% humidity, 20°C day temperature, 6°C night temperature, 16 h/8 h light/dark cycle and 250 µmol m-2 s-1 light intensity. Seedlings were then transferred to a virus nursery and singled out into 6-cm pots at 60% humidity, 20°C/16°C at 11.5 h/12.5 h light/dark cycle and 120 μE. After infection, plants were placed in a quarantine chamber under long-day conditions (70% humidity, 23°C/20°C at 16 h/8 h light/dark cycle and 120 μE) to promote flowering.

### Virus inoculation

TYMV-S strain was isolated in the Czech Republic from Chinese cabbage and is commonly used in laboratories and companies (BioReba). TYMV-M strain was obtained from DSMZ (order number PV0299) as desiccated inoculum (genbank ID: ON924209). *A. thaliana* Col-0 plants were infected with primary inoculum and a new inoculum stock used throughout this study was generated by harvesting and shock-freezing systemically infected Arabidopsis leaves. For infection experiments, frozen inoculum stock was ground with mortar and pestle in 50 mM sodium phosphate buffer, pH 7.2 supplemented with 0.2% sodium sulfite. Celite 545 (Merck) was added to inoculum before rub infection with cotton swabs of four to six fully extended leaves.

### Crosses and vertical transmission assays

For all crosses, designated female flower buds were emasculated and the pistils were hand-pollinated two days after emasculation. For pollination from infected plants, open flowers from symptomatic branches with visible pollen from at least three individual plants were used as pollen donors for each experiment. For vertical transmission counting, dry seeds were sown directly on soil. Seeds were vernalized at 4°C for 2-4 days before transfer to a growth chamber with long day conditions. Vertical transmission rate was assessed visually between 10 and 21 days after germination. All counted seedlings were removed and the procedure repeated until no seedlings remained in soil.

### Pollen isolation

Pollen was harvested approximately four weeks after virus infection by submerging open flowers from 10 – 60 plants directly in 0.3 M ice-cold mannitol. Samples were vigorously vortexed to release the pollen and filtered through a 50 µm mesh. The mesh was subsequently washed with an additional 1 ml of 0.3 M mannitol. Pollen was spun down at 3000 g for 3 min at 4°C and the supernatant discarded. Samples were resuspended in 500 µl cold 0.3 M mannitol substituted with 0.1% Triton X100 and 1 U Ribolock. Samples were again filtered through a 50 µm mesh and spun down at 3000 g for 3 min at 4°C. Supernatant was removed to leave about 100 µl pollen suspension which was either flash-frozen in liquid nitrogen until RNA extraction followed by qRT-PCR or sRNA sequencing or was directly used for FISH. Open flowers were directly submerged into 4% PFA for *qrt1-4* pollen harvest for subsequent FISH. Samples were quickly vortexed to release pollen and the pollen let settle by gravity without additional filtering before continuing with FISH protocol.

### Molecular cloning

All binary plasmids used in this work were generated through Golden Gate cloning, using the GreenGate modular system (*47*), as previously described (*20*). The entry vectors were used to assemble into pGGSun (*20*) the following cassettes: *pAGO5::MYC:AGO5, pHTR10::AGO5, pWOX2::MYC:AGO5, pHTR10::siTYMV, pUBI10::siTYMV, pWOX2::siTYMV, pHTR10::siScar, pUBI10::siScar, pWOX2::siScar* and *pWOX2::H2B:Clover*. In pGGZ (*47*) the following cassettes were assembled: *pHTR10::P19* and *pHTR10::H2B:Clover*.

### *In situ* hybridization (ISH)

*In situ* hybridization was performed as described in (*20*). In short, dissected inflorescence tissues of interest were stained in 1% w/v eosin in 70% ethanol, then infiltrated with xylene substitute and paraffin, then cast into paraffin blocks. These were then cut into 2 μm-thick sections that were transferred onto glass microscopy slides and screened for areas of interest. DIG-labeled RNA probes to detect TYMV were generated with the DIG RNA Labeling Kit T7/SP6 (Roche #11175025910); see (*20*) for primers used to generate the DNA templates. Slides were twice incubated 10 min in Histo-Clear II (National Diagnostics #HS-202), twice 5 min in 100% ethanol, then rehydrated through serial passages in 90%, 70%, 50%, and 30% ethanol, then in Tris-EDTA pH 7.5. Sections were then treated with Proteinase K (Roche #3115836001), washed in 1× PBS, then incubated for 10 min in 4% paraformaldehyde, dehydrated through serial ethanol washes, and air-dried. After probe denaturation for 3 min at 80°C, hybridization with 50-100 ng DIG-labeled probes per slide was performed O/N at 50°C in 150 μl hybridization solution: 50% formamide, 10 mM Tris base, 300 mM NaCl, 5 mM EDTA, 10 mM Na2HPO^4^, 1× Denhardt’s solution (Sigma Aldrich #D2532-5ML), 10% dextran sulfate, and 0.5 μg/μl tRNA (Roche #10109517001). Slides were washed in 2× SSC, then incubated in 0.2× SSC for 2 h at 55°C and treated with RNase A (Thermo Scientific #EN0531) at 37°C for 30 min, then 1 h in 0.2× SSC at 55°C. Slides were washed 10 min in washing buffer and incubated 1 h in blocking buffer (Roche #11585762001). Anti-DIG antibody was added (Roche #11093274910 - 1:1500 dilution) and incubated 1 h 45 min at room temperature, washed for 1 h, incubated in TNM5 (100 mM Tris pH 9.5, 100 mM NaCl, 5 mM MgCl^2^) three times for 2 min, then O/N in TNM5 with 10% w/v polyvinyl alcohol, and 10 μl/ml NBT/BCIP (Roche #11697471001). Slides were mounted with Aqua-Poly/Mount (Polysciences #18606-20) and scanned at 40× magnification.

### Fluorescent *in situ* hybridization (FISH)

Whole-mount fluorescent *in situ* hybridization was performed as previously described (*32*), with some modifications. Harvested tissues of interest were placed in fixation buffer (4% paraformaldehyde in 1X PBS) and vacuum-infiltrated, then incubated at room temperature for a minimum of 30 min and washed twice with 1X PBS. To digest cell walls, the tissues were incubated in 2% w/v cellulase (Serva #16419.03) and 1% w/v macerozyme (Serva #28302.02) in PBS for 30 min with gentle shaking, then washed three times in 1X PBS. Subsequently, tissues were dehydrated through two 15-min incubations in 100% methanol and one in 100% ethanol, then incubated in ClearSee Alpha (xylitol 10% w/v, sodium deoxycholate 15% w/v, urea 25% w/v in water with 50 mM sodium sulfite (*48*)) for different periods of time depending on the tissue (meristems ≥ 3 days; pollen ≥ 3 days; carpels ≥ 20 days; fertilized ovules ≥ 28 days). Tissues were washed twice for 15 min with 1X Stellaris RNA FISH Wash Buffer A (Biosearch Technologies, #SMF-WA1-60) supplemented with 10% (v/v) formamide (Millipore, #S4117), then arranged on poly-L-lysine-coated slides (Thermo Scientific, #10219280) and allowed to air-dry. 100 µl embedding solution (10% acrylamide:bisacrylamide) was applied to the samples and covered with a coverslip, then polymerized by adding APS and TEMED and leaving at room temperature for 30 min. After removing the coverslip, 100 µl hybridization solution was added (100 mg/ml dextran sulfate (Sigma, #D8906-10G), 10% formamide, 2X SSC buffer (Invitrogen, #AM9763), probe concentration of 250 nM). After adding a coverslip, the slide was placed in a humid box overnight at 37°C. The coverslip was carefully removed, the slide was placed in a Coplin jar wrapped with aluminum foil and submerged in 10% formamide, 2X SSC in RNAse-free water, twice for 30 min. After removing excess buffer, 80–100 µl of Vectashield Antifade Mounting Medium with DAPI (BioZol #VEC-H-1200) were added, the sample covered with a coverslip and sealed with nail varnish. Samples were imaged using Leica Stellaris 8 or Leica SP8 laser confocal microscopes, with 20X and 40X water-immersion objectives. Quasar570 probe fluorescence was detected through excitation at 561 nm laser and emission between 567-599 nm. DAPI was detected at 405 nm excitation, emission between 415-490 nm. Images were then analyzed and processed with FIJI. Stellaris probe sets of DNA oligonucleotides (**Table S3**) labelled with the Quasar570® fluorophore were designed with Stellaris RNA FISH Probe Designer (www.biosearchtech.com/stellarisdesigner) and synthesized by BioCat.

### RNA blotting and RT-qPCR

RNA was extracted using TRI reagent (Zymo Research #R2050-1-200). In short, harvested plant tissues were flash-frozen in liquid nitrogen and either ground with pestle and mortar or metal beads before addition of TRI reagent. After removing debris by centrifugation, chloroform was used to separate aqueous and organic phase. Isopropanol was subsequently added to the aqueous phase and incubated for 2 h at -20°C to precipitate RNA. RNA was pelleted through centrifugation and washed twice with cold 80% ethanol. The resulting RNA pellet was dried and resuspended in RNase-free water and concentration and quality were measured. RNA was stored at -80°C until further use. Northern blotting to detect viral RNA was performed as previously described (*20*) . Briefly, 5 μg RNA was denatured by incubating with 15% v/v deionized glyoxal at 50°C for 1 h. Samples were run in 1% agarose gel in 20 mM sodium phosphate pH 7.2. Capillary transfer to nylon membrane was performed overnight, after which RNA was crosslinked to the membrane by UV irradiation. After staining with methylene blue, membranes were probed with γ-^32^P-ATP-labeled @TYMV DNA oligonucleotides, described in (*20*).

RT-qPCR was performed by first treating the RNA with DNAse (Thermo Fischer Scientific TURBO DNA-free kit, #AM1907), then performing random hexamer-primed reverse transcription on 500ng RNA (Thermo Fischer Scientific RevertAid First Strand cDNA Synthesis kit, #K1622). qPCR was performed on 1:40 diluted cDNA (Applied Biosystems PowerSYBR Green PCR Mastermix, #4367659). UBC28 ([AT1G64230](https://www.arabidopsis.org/access-denied)) (*49*) and PP2A (AT1G69960) (*50*) were used as references genes (Table S1 for primers).

### Protein extraction, immunoblotting and ELISA

Proteins for immunoblot analysis were isolated by homogenizing frozen plant tissue in extraction buffer (0.7 M sucrose, 0.5 M Tris-HCl pH 8, 5 mM EDTA pH 8, 0.1 M NaCl), mixing the same volume of the homogenate with Tris-EDTA-equilibrated phenol pH 8 8, centrifugating and transferring the phenolic phase to a new tube with 4 volumes 0.1 M ammonium acetate dissolved in methanol, then incubating overnight at -20°C. After centrifugation, the pellet was washed twice in 0.1 M ammonium acetate dissolved in methanol, dried and resuspended in 10% glycerol, 3% SDS, 62.3 mM Tris-HCl pH 8. After protein quantification by amido black, Laemmli buffer was added and samples were stored at -20°C. For western blot analysis, samples were first heated at 70°C for 10 min, then loaded into precast gradient 4-20% MP TGX Stain-Free Gel (BioRad, #4568096). Proteins were then transferred onto PVDF membrane (Merck Immobilon-P, #IPVH00010) in Tris-glycine/20% ethanol, then blocked in TBS-Tween 5% skim milk powder for at least 1 h before adding HRP-coupled c-myc antibody (Myltenyi #130-092-113) and incubating overnight at 4°C. The membrane was then washed three times in TBS-Tween and chemiluminescence revealed through SuperSignal West Atto substrate (Thermo Fischer Scientific #A38555). The membrane was then stained with amido black (10% acetic acid, 0.1% naphthol blue black).

To detect TYMV in seedlings, we used ELISA according to manufacturer’s protocol (https://www.bioreba.ch/product/TYMV-Complete-kit-9601.aspx).

### Small RNA sequencing and analysis

Small RNA library preparation and sequencing was performed by Novogene. Prior to library preparation, RNA quality was evaluated with Agilent Bioanalyzer 5400/AATI. Initial size selection from total RNA was performed by isolating 14-40 bp fragments from an agarose gel. 3’ and 5’ adaptors were ligated to the ends of small RNA, then the first strand cDNA was synthesized. Double-stranded cDNA libraries were generated through PCR enrichment. After purification and size selection, libraries were checked with Qubit and real-time PCR for quantification and bioanalyzer for size distribution. Libraries were sequenced with a Novaseq 6000 sequencing system, approximately 20 M reads per sample, single-end 50 bp (SE50).

Raw reads were first trimmed using with Trimgalore (version 0.6.10) with adjusted parameters (-j 8 -q 30 --length 20 --max_length 25 –fastqc). Trimmed reads were then mapped against the siScarlet construct, the siTYMV construct using Bowtie-1.3.1 (settings: -q -v 0 -k 10 -S -t --threads 8). Files were then converted into BAM files and sorted and indexed using Samtools-1.19.2. Size distributions of mapped reads were generated using the samtools view in combination with: awk ‘print length($10)}’ | sort | uniq -c).

### Data analysis and statistical methods

Data were visualized using R Studio and packages “dyplr” (*51*) and “ggplot2” (*52*) and figure panels assembled in Affinity Designer2 version 2.6.3. Differences in vertical transmission after selfing were analyzed using One-way analysis of variance (ANOVA) followed by a post-hoc Tukey HSD test (α = 0.05) was performed with R version 4.3.1 and the R-package “agricolae” (version 1.3-3; https://cran.rproject.org/web/packages/agricolae/index.html). Differences in pollen transmission between genotypes were analyzed using a generalized linear model (GLM) with a quasibinomial distribution and a logit link function with R version 4.3.1 and the packages “emmeans” (*53*), “multcomp” (*54*), “lme4” (*55*). See Table S1 for statistical analyses.

