## Supplemental Figures S1 to S13 for "AGO5 restricts virus vertical transmission in plant gametophytes"

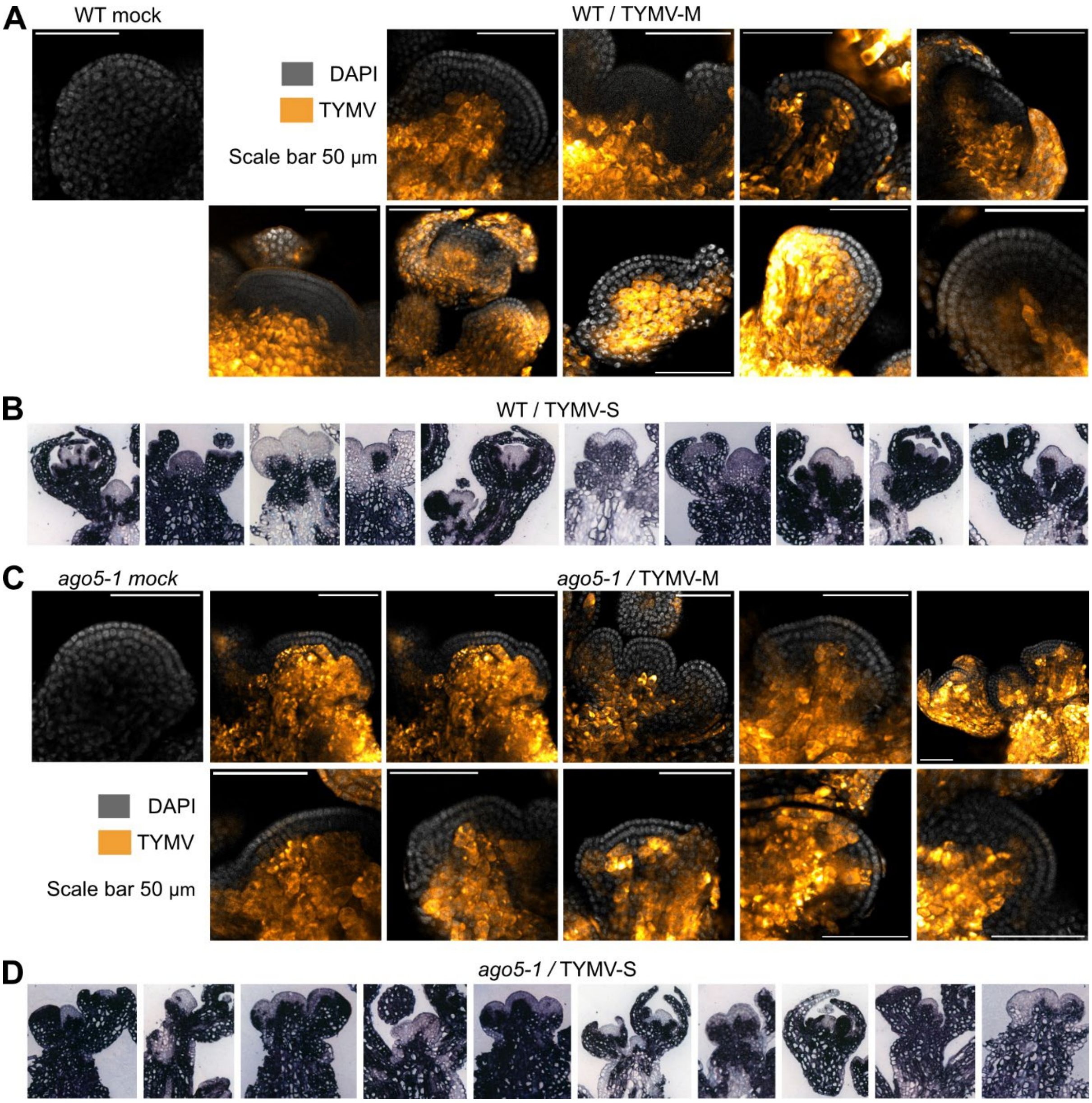


**Figure S1: TYMV is excluded from shoot apical and flower meristem in WT and *ago5* plants.** Supports Fig. 1A. **(A,C)** Additional images of FISH detection of TYMV-M positive sense RNA in Arabidopsis WT (A) and *ago5-1* (C) shoot apical and flower meristems. Images show merge of TYMV FISH channel (pseudo-colored in LUT „orange hot") and DAPI (greyscale). Scale bars = 50 µm. **(B,D)** *In situ* hybridizations on vertical sections of WT (B) and *ago5-1* (D) meristems with a DIG-labelled probe to detect TYMV-S. Hybridization of probe is visible in dark purple.

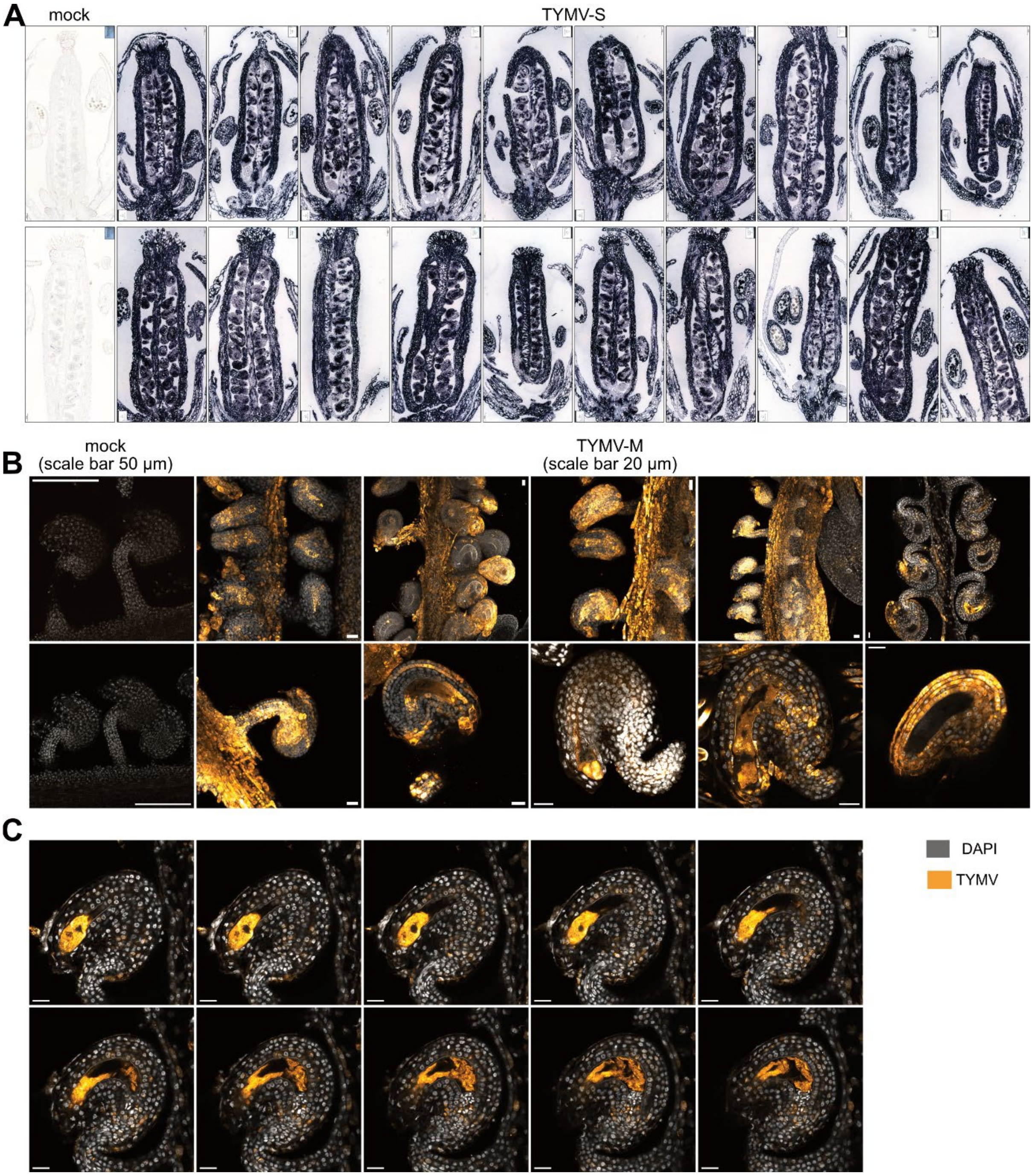


**Figure S2: TYMV can infect all parts of Arabidopsis gynoecia.** Supports Fig. 1B,C. (**A)** *In situ* hybridizations on vertical sections of open flowers from non-infected (far left) and TYMV-S-infected plants, with a DIG-labelled probe to detect TYMV. Hybridization of probe is visible in dark purple. **(B)** Additional images of FISH detection of TYMV-M genomic RNA in Arabidopsis ovules. The overview top images are max projections. **(C)** Z-series in 2µm increments through ovule shown in Figure 1B. In B and C images show merge of TYMV FISH channel (pseudo-colored in LUT „Orange Hot") and DAPI (greyscale). Scale bars = 50 µm (mock) or 20 µm (TYMV-M infected).

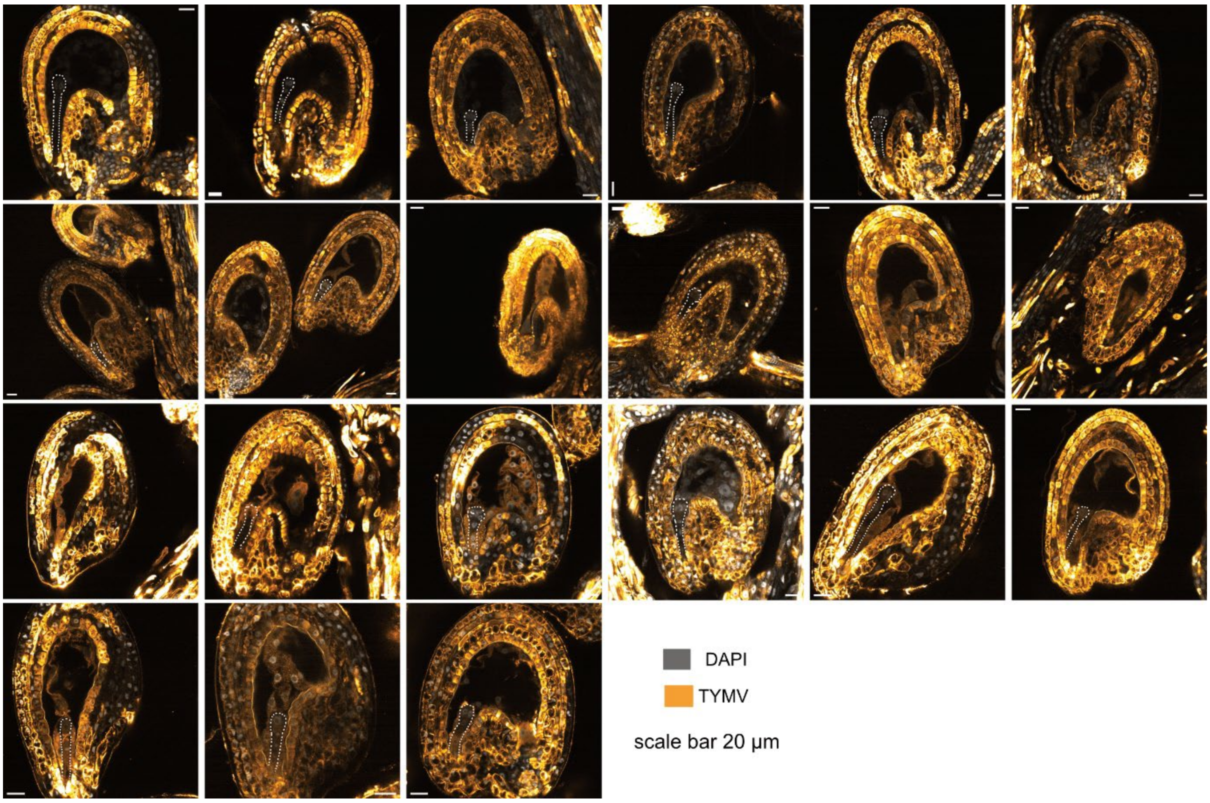


**Figure S3: Maternal transmission of TYMV-M in endosperm and embryo 48 hours after pollination.** Supports Figure 1D-F. Additional laser confocal microscopy images of TYMV-M genomic RNA detection by FISH in Arabidopsis seeds, 48 hours after pollination with pollen from non-infected plants. Images show merge of TYMV FISH channel (pseudo-colored in LUT „Orange Hot") and DAPI (greyscale). Scale bars = 20 µm. Embryos are outlined by dotted line for clarity.

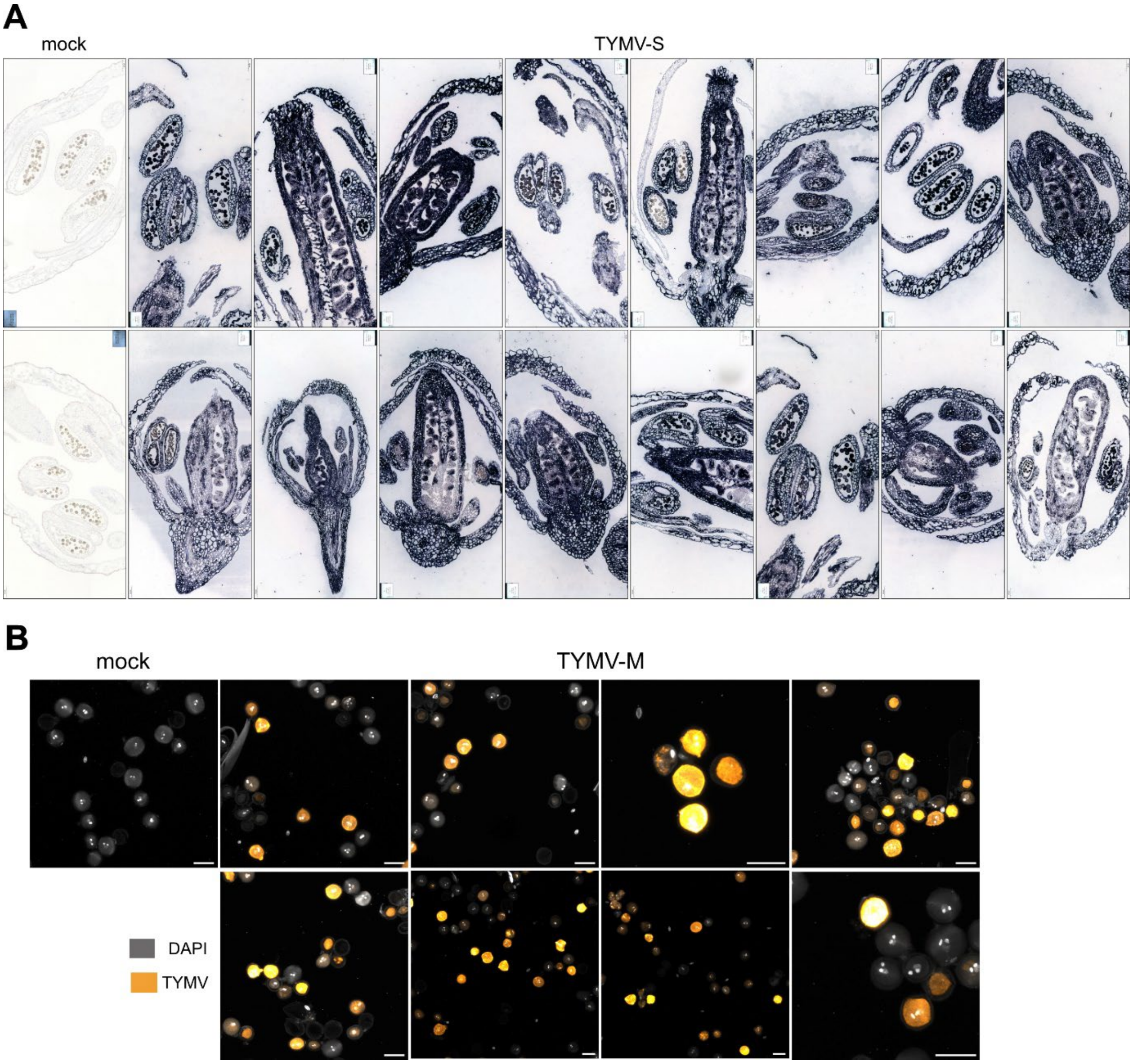


**Figure S4: Detection of TYMV in anthers and pollen.** Supports Fig. 1G,H. **(A)** *In situ* hybridizations on vertical sections of open flowers from non-infected (far left) and TYMV-S-infected plants, with a DIG-labelled probe to detect TYMV RNA. Hybridization of probe is visible in dark purple. **(B)** Additional laser confocal microscopy images of TYMV-M genomic RNA detection by FISH in Arabidopsis pollen. Images show merge of TYMV FISH channel (pseudo-colored in LUT „Orange Hot") and DAPI (greyscale). Scale bars = 20 µm.

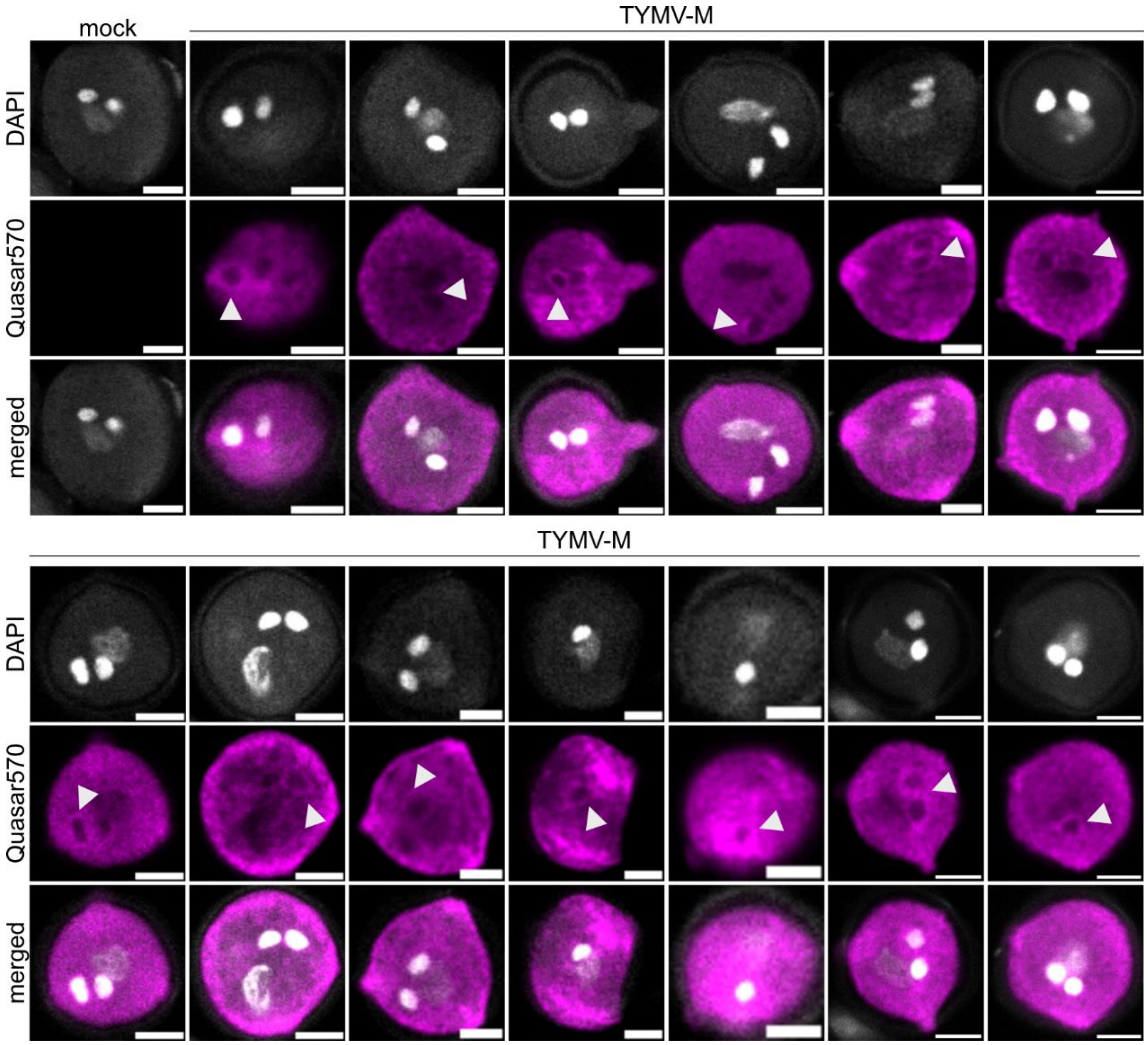


**Figure S5: Detection of TYMV-M in sperm cell cytoplasm of mature pollen.** Supports Fig. 1H. Close-up laser confocal images of single pollen grains showing TYMV RNA detection by FISH in sperm cell cytoplasm (arrowheads) and nuclear exclusion. Images show sperm cell nuclei and vegetative nuclei stained by DAPI (grey, upper row), TYMV FISH channel (magenta, middle row) and merged images (lower row). Scale bars = 5 µm.

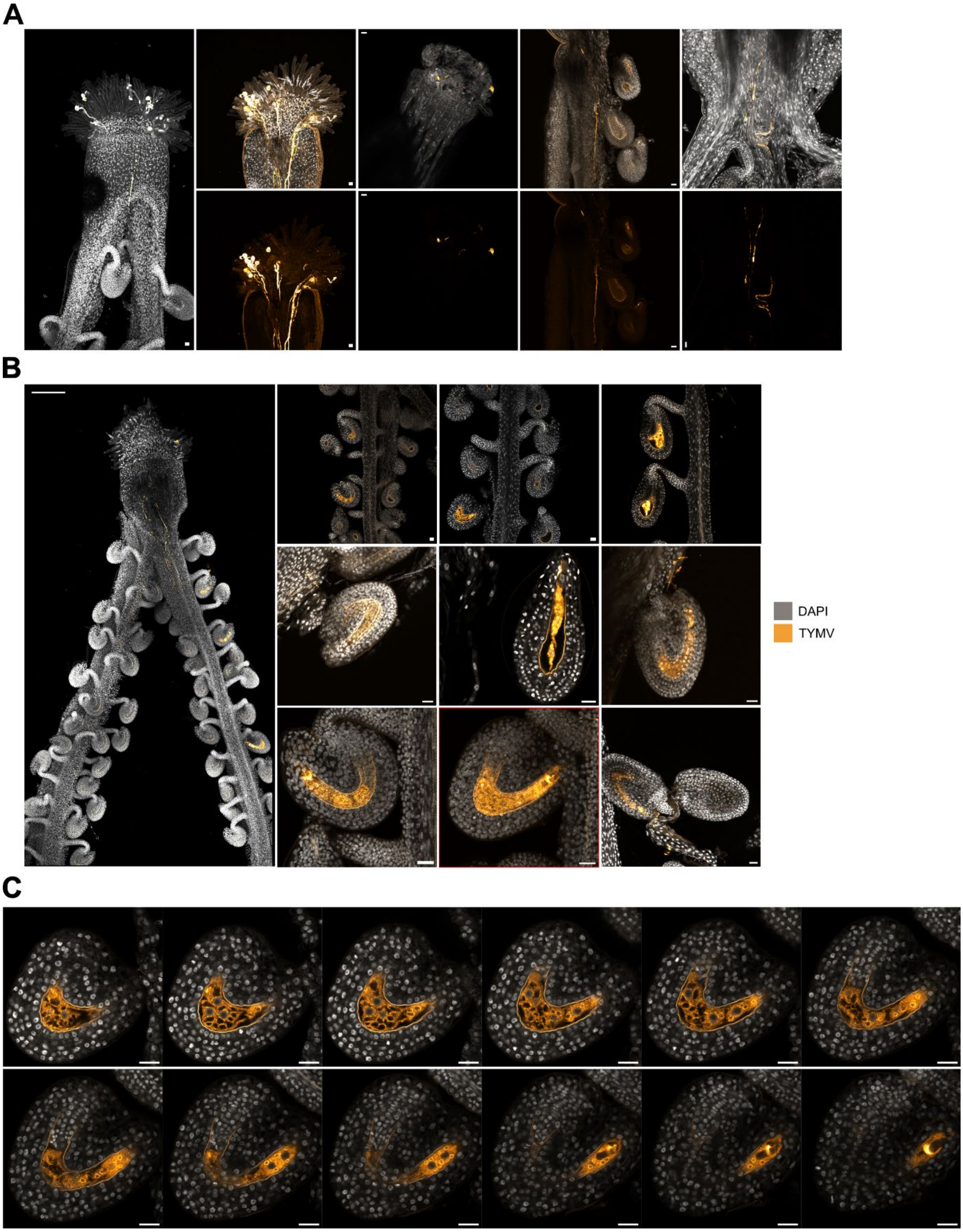


**Figure S6: Transmission via pollen of TYMV-M**. Supports Fig. 1I,J. **(A)** Additional laser confocal microscopy images (max projections) of TYMV genomic RNA detection by FISH in pollen tubes growing into healthy pistils 24 hours after pollination. Top images show merge of TYMV FISH (pseudo-colored in LUT "Orange Hot") and DAPI (greyscale) channels, bottom images show only FISH signal. Scale bars = 20 µm. **(B)** Additional laser confocal microscopy images (mostly max projections) of TYMV-M genomic RNA detection by FISH in fertilized endosperm 24 hours after pollination. A **(C)** Z-series in 2µm increments through the endosperm cavity shown in Figure 1J. Images show merge of TYMV FISH (pseudo-colored in LUT "Orange Hot") and DAPI (greyscale) channels. Scale bars = 20 µm, 200µm for images on the left in A and B.

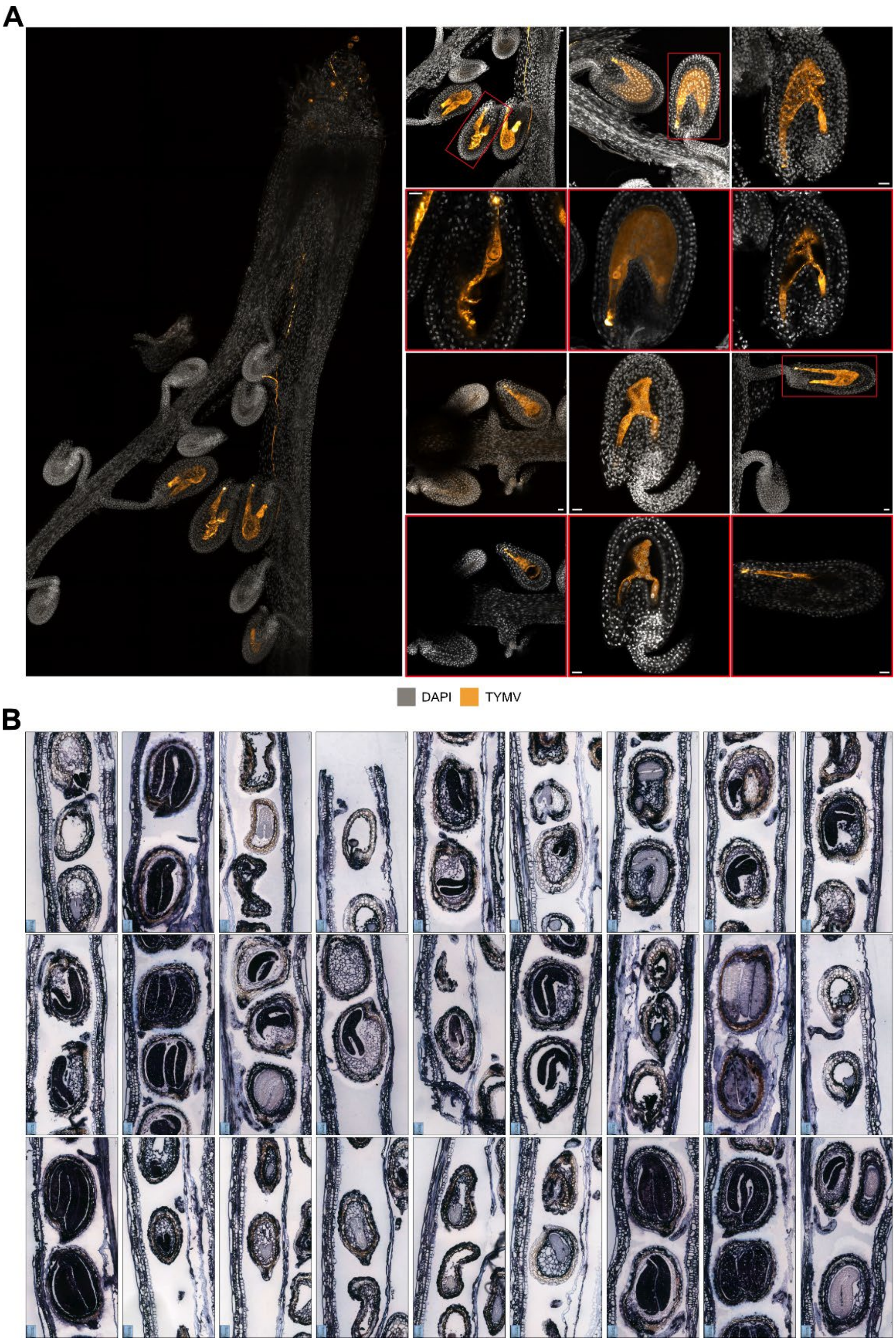


**Figure S7: Embryo infection by TYMV after transmission via pollen.** Supports Fig. 1K,L. **(A)** Additional laser confocal microscopy images of TYMV-M genomic RNA detection by FISH in Arabidopsis seeds 48 hours after pollination of non-infected mother with pollen from TYMV-infected father. Red-boxed images are single frames of Z-stacks focusing on the embryo. Images show merge of TYMV FISH channel (pseudo-colored in LUT "Orange Hot") and DAPI (greyscale). Scale bars = 20 µm. **(B)** *In situ* hybridizations on vertical sections of developing Arabidopsis siliques from TYMV-S-infected plants, with a DIG-labelled probe to detect TYMV RNA. Hybridization of probe is visible in dark purple.

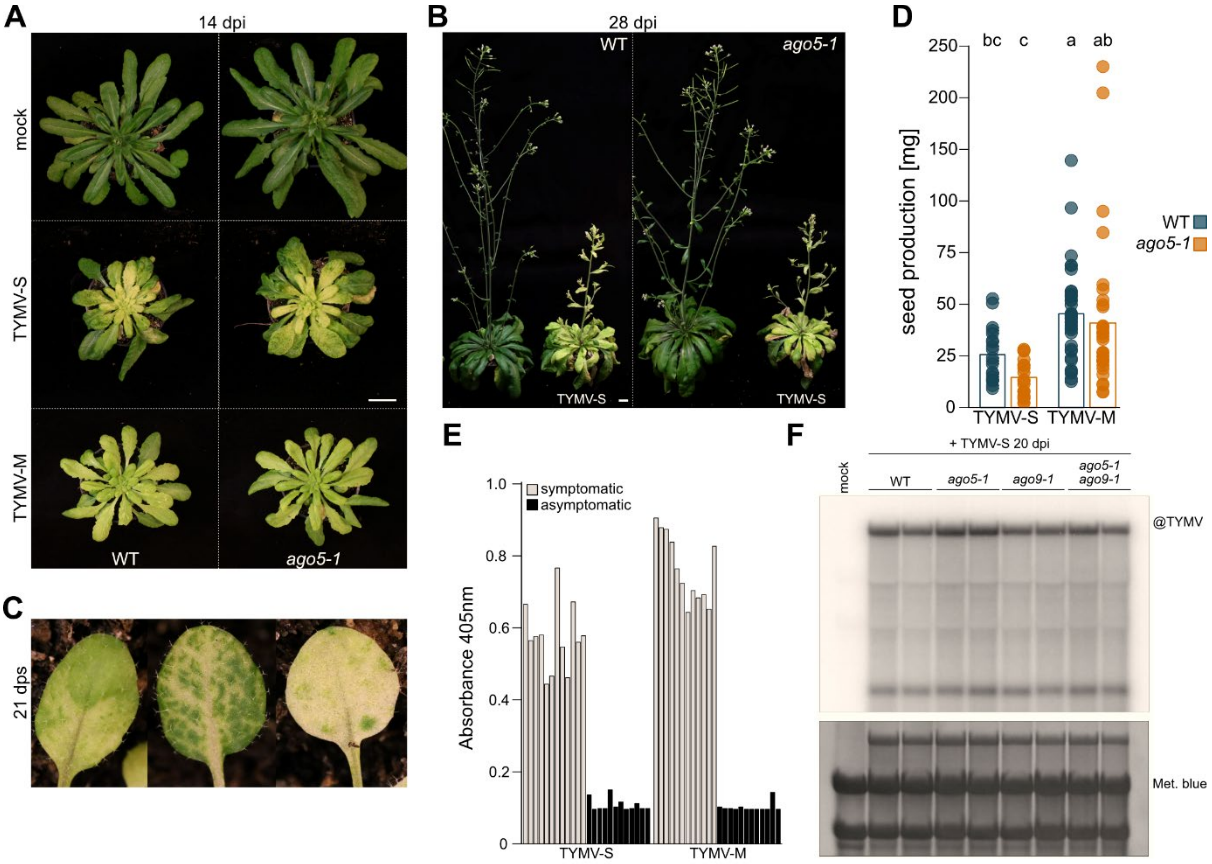


**Figure S8: Infection traits of TYMV-S and -M isolates in Arabidopsis.** Supports Fig. 2. **(A)** Photographs of systemically infected WT and *ago5-1* plants 14 days after rub-inoculation with mock buffer (top) or sap from TYMV-S (middle) or TYMV-M (bottom) infected plants. Scale-bar: 2 cm. **(B)** Photographs of flowering WT and *ago5-1* TYMV-S-infected plants at 28 dpi. Scale-bar: 2 cm. **(C)** Photographs detailing TYMV-S symptoms in infected progeny of systemically infected WT plants, following vertical transmission, displaying different degrees of yellowing in the first true leaves at 21 days post-sowing (dps). **(D)** Seed production (mg per plant) by TYMV-infected WT and *ago5-1* plants. Dots represent individual plants, vertical bars the average. Seed weights correspond to plants used in Fig. 2B for TYMV-S and Fig. 2C, 3F and S11B for TYMV-M. **(E)** ELISA absolute absorbance at 405nm, to detect TYMV in progeny of systemically infected plants, following vertical transmission, at 18 days post-sowing. **(F)** Northern blot analysis of TYMV-S RNA accumulation in inflorescences of WT and *ago5-1*, *ago9-1* and *ago5-1/ago9-1* plants, each in duplicate of two pools of 4-5 plants at 20 dpi. Methylene blue staining is used as loading control.

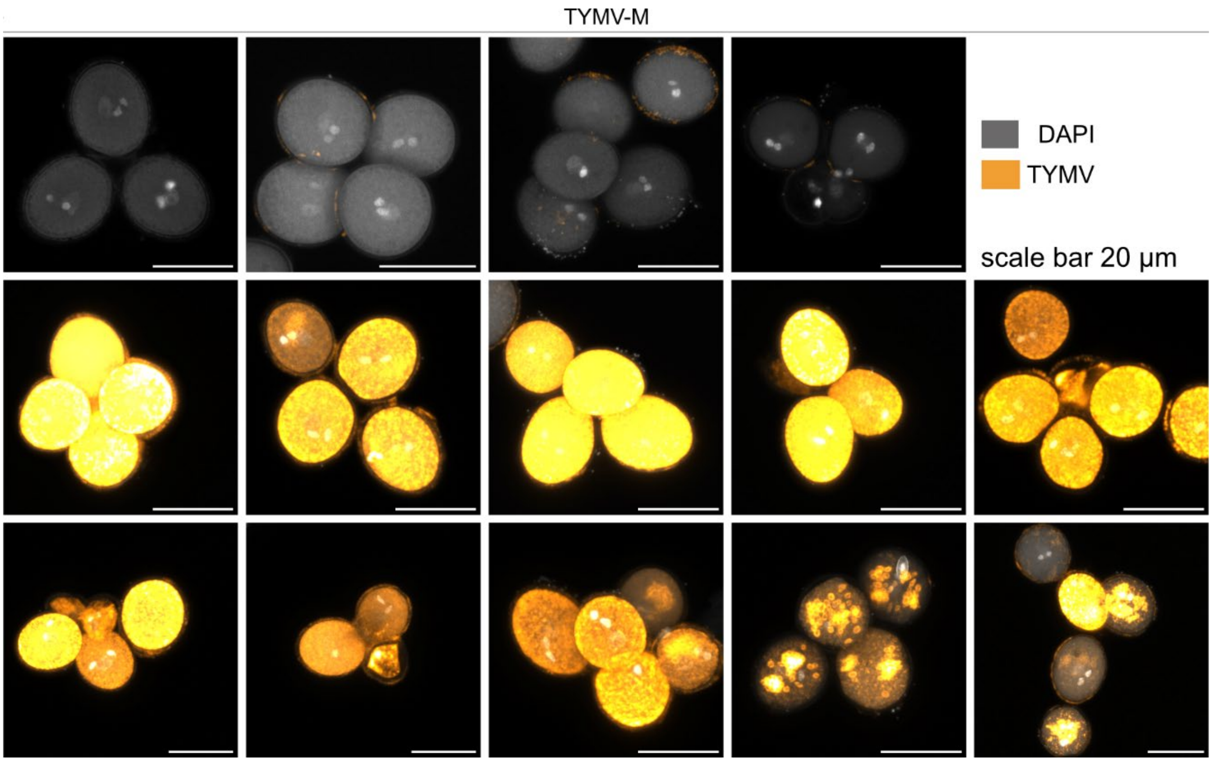


**Figure S9: TYMV-M infected pollen tetrads of *qrt1-4*.** Supports Fig. 3. Additional laser confocal microscopy images (max projections) of TYMV-M genomic RNA detection by FISH in *qrt1-4* pollen tetrads. Images show merge of TYMV FISH (pseudo-colored in LUT “Orange Hot”) and DAPI (greyscale) channels. Scale bars: 20 µm.

**A**

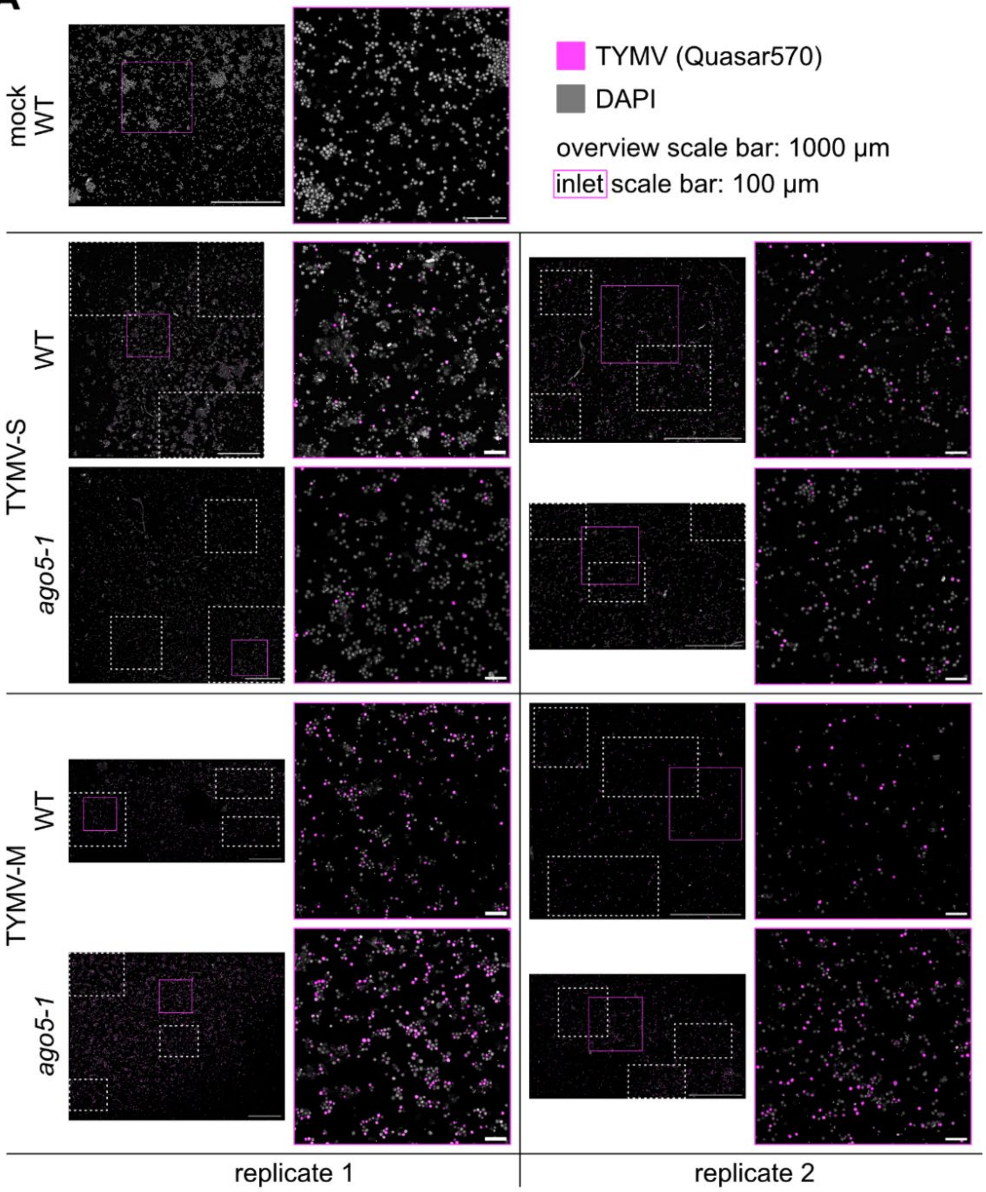


**B**

| | | | replicate 1 | | replicate 2 | |
|---|---|---|---|---|---|---|
| | | | DAPI | TYMV | DAPI | TYMV |
| TYMV-S | WT | R1 | 866 | 180 | 198 | 46 |
| | | R2 | 254 | 40 | 127 | 29 |
| | | R3 | 488 | 95 | 89 | 27 |
| | ago5-1 | R1 | 504 | 62 | 98 | 25 |
| | | R2 | 1291 | 189 | 155 | 37 |
| | | R3 | 355 | 50 | 135 | 37 |
| TYMV-M | WT | R1 | 131 | 116 | 43 | 25 |
| | | R2 | 388 | 300 | 99 | 68 |
| | | R3 | 257 | 208 | 94 | 51 |
| | ago5-1 | R1 | 201 | 249 | 98 | 87 |
| | | R2 | 173 | 186 | 81 | 65 |
| | | R3 | 525 | 519 | 107 | 95 |

**Figure S10: TYMV-infected pollen grain quantification.** Supports Fig. 3. **(A)** Laser confocal microscopy images (max projections) of TYMV genomic RNA detection by FISH for quantification of presence/absence of TYMV-S and -M in mature pollen grains. Images show merge of TYMV FISH (magenta) and DAPI (greyscale) channels. The first image of each quadrant shows stitched overview image used for counting. Counted areas (R1-2-3) are marked by white boxes. Inlets for better visualization are marked by magenta box outlines. Scale bars: 1000 µm for overviews and 100 µm for inlets. **(B)** Absolute counts of TYMV presence/absence in Arabidopsis pollen grains corresponding to images shown in (A).

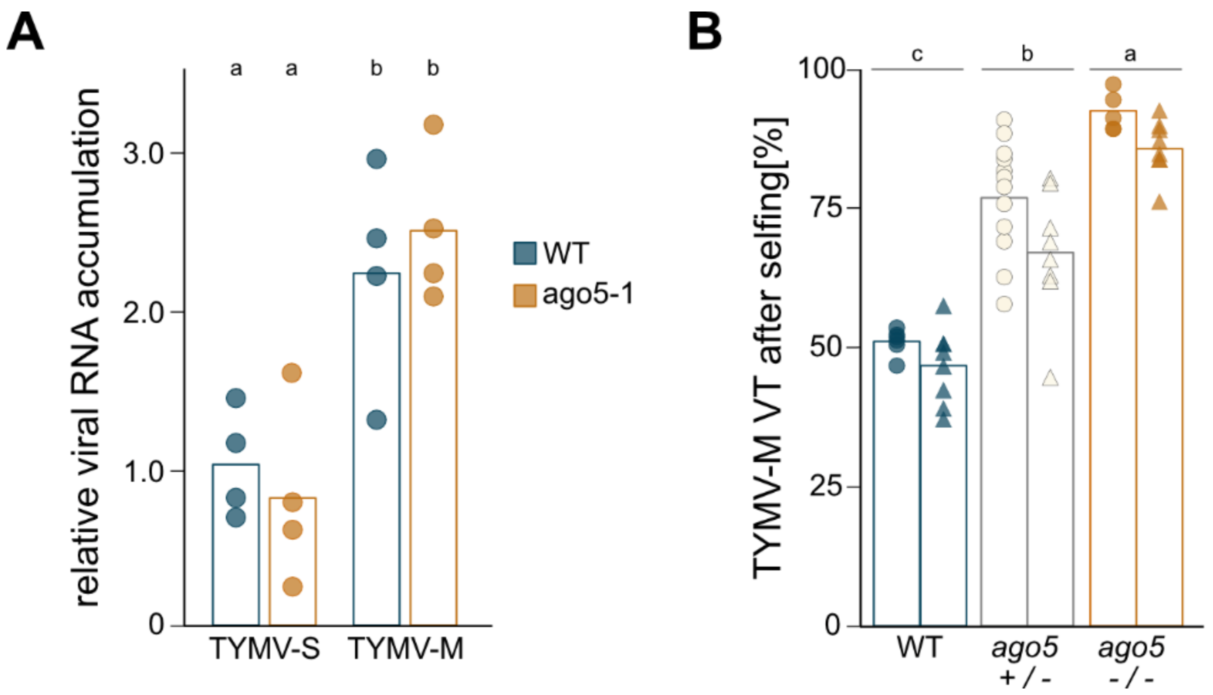


**Figure S11: TYMV accumulation and transmission.** Supports Fig. 3. **(A)** qRT-PCR quantification of TYMV-S and TYMV-M RNA in mature pollen in WT and *ago5-1*. **(B)** VT rates [%] of TYMV-M after selfing in indicated genotypes. Single data points refer to individual parent plants, shape of data points to independent infection experiments. Seedlings counted (sc): 16113. Statistical significance was determined by one-way ANOVA coupled with Tukey's HSD test ($\alpha = 0.05$), lower case letters indicate statistical groups.

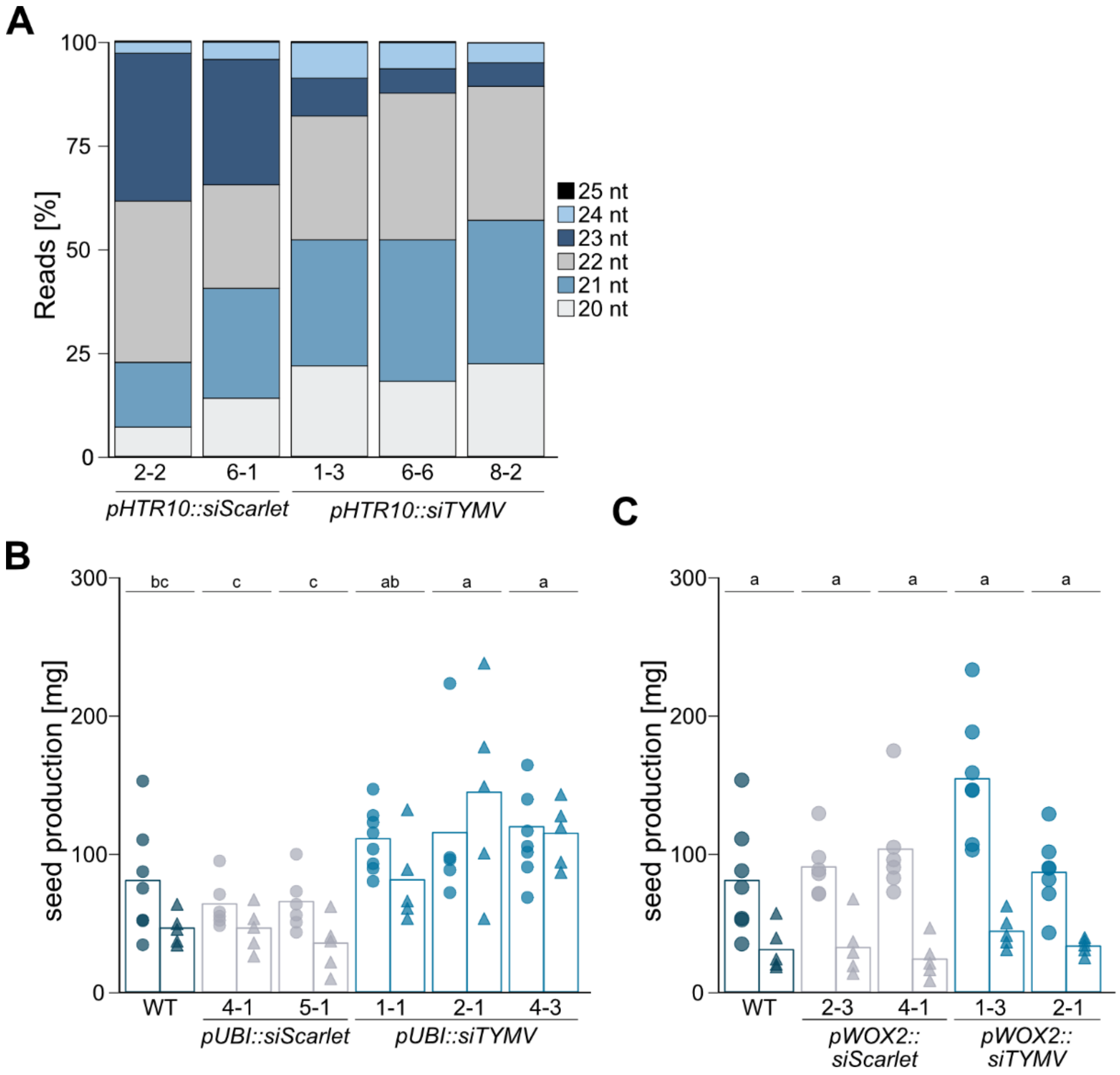


**Figure S12: RNA hairpin-mediated production of vsiRNA throughout reproduction**. Supports Fig. 4. **(A)** Read length distribution of 20-25 nt siRNA in *pHTR10::siTYMV/siScarlet*/WT transgenic lines. **(B)** Seed production (mg per plant) by TYMV-M-infected *pUBI::siTYMV/siScarlet*/WT plants, corresponding to VT rates displayed in Fig. 4C. Single data points refer to individual parent plants, shape of data points to independent infection experiments. **(C)** As in (B), but by *pWOX2::siTYMV/siScarlet*/WT

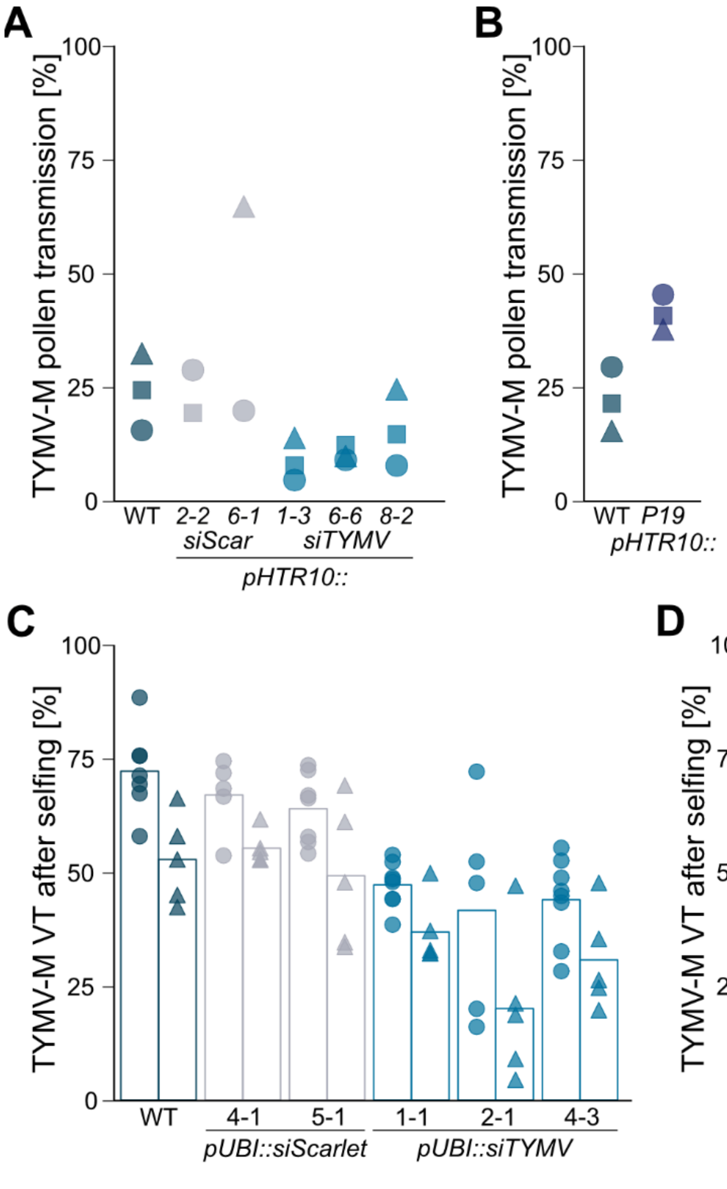


**Figure S13: TYMV VT in hairpin-expressing transgenic lines.** Supports Fig. 4. **(A)** Absolute VT via pollen [%] of TYMV-M in *pHTR10::siScarlet*/WT and *pHTR10::siTYMV*/WT, corresponding to relative data shown in Fig. 4B. **(B)** As in (A), but in *pHTR10::P19*/WT. Single data points refer to individual parent plants, shape of data points to independent infection experiments. **(C)** Absolute VT rates [%] of TYMV-M after selfing of *pUBI::siTYMV/siScarlet*/WT lines, relative rates shown in Fig. 4C. **(D)** As in (C), but of *pWOX2::siTYMV/siScarlet*/WT lines, relative data shown in Fig. 4D.
